# Deciphering the involvement of the Hippo pathway co-regulators, YAP/TAZ in invadopodia formation and matrix degradation

**DOI:** 10.1101/2022.06.28.497902

**Authors:** Jubina Balan Venghateri, Bareket Dassa, David Morgenstern, Michal Shreberk-Shaked, Moshe Oren, Benjamin Geiger

## Abstract

Invadopodia are adhesive, actin-rich protrusions, formed by metastatic cancer cells that degrade the extracellular matrix and facilitate invasion. They support the metastatic cascade by a spatially and temporally coordinated process whereby invading cells bind to the matrix, degrade it by specific metalloproteinases, and mechanically penetrate diverse tissue barriers by forming actin-rich extensions. However, despite the apparent involvement of invadopodia in the metastatic process, the molecular mechanisms that regulate invadopodia formation and function are still largely unclear. In this study, we have explored the involvement of the key Hippo pathway co-regulators, namely YAP, and TAZ, in invadopodia formation and matrix degradation. Towards that goal, we tested the effect of depletion of YAP, TAZ, or both on invadopodia formation and activity in multiple human cancer cell lines. We report that knockdown of YAP and TAZ or their inhibition by verteporfin induce a significant elevation in matrix degradation and invadopodia formation in several cancer cell lines. Conversely, overexpression of these proteins strongly suppresses invadopodia formation and matrix degradation. Proteomic and transcriptomic profiling of MDA-MB-231 cells, following co-knockdown of YAP and TAZ, revealed a significant change in the levels of key invadopodia-associated proteins, including the crucial proteins Tks5 and MT1-MMP (MMP14). Collectively, our findings show that YAP and TAZ act as negative regulators of invadopodia formation in diverse cancer lines, most likely by reducing the levels of essential invadopodia components. Dissecting the molecular mechanisms of invadopodia formation in cancer invasion may eventually reveal novel targets for therapeutic applications against invasive cancer.

## Introduction

Tumor metastasis is a complex multi-step process that accounts for the vast majority of all cancer-related deaths (1,2). The metastatic invasion process is commonly initiated by the loss of intercellular cohesion in the primary tumor, followed by an invasion of the cancer cells, individually or collectively, into the surrounding tissues (3). The invasive phase of the metastatic process, whereby the cancer cells penetrate into the nearby extracellular matrix (ECM) and blood vessels, utilizes tightly-coordinated adhesive-proteolytic-protrusive processes, combining integrin-mediated adhesion to the ECM, secretion of matrix metalloproteinases, and activation of cytoskeleton-based protrusive machinery that physically penetrate the surrounding connective tissue (4). Throughout this process, the metastatic cancer cells sense the properties of the ECM and respond to the chemical and physical cues that eventually guide them to the neighboring vasculature (4).

The mechanical pressure applied by invading cells to the nearby matrix is generated by different membrane protrusions (e.g., filopodia, lamellipodia, ruffles, and invadopodia), the most prominent of which are invadopodia that contain an integrin-based ECM adhesion domain, a protrusive actin-rich core, and diverse proteinases that degrade the matrix (5–8). Invadopodia were first identified over 30 years ago in embryonic fibroblasts, transformed with Rous Sarcoma Virus (6,9,10), and later observed in a wide variety of cancers, including melanoma, head and neck tumors, as well as breast, pancreatic, and prostate carcinomas, to list just a few (11,12). Notably, some non-cancerous cells, such as dendritic cells, macrophages, endothelial cells, vascular smooth muscle cells, and osteoclasts, possess similar adhesion structures that are believed to partake in their physiological migratory and tissue remodeling activities (6,13).

The invadopodia structural core comprises of F-actin and several cytoskeleton-modulating and scaffolding proteins (e.g., cortactin, Tks4, Tks5, Wiskott-Aldrich Syndrome Protein (N-WASP), specific adhesion proteins (mainly integrins), as well as diverse signaling molecules, and their regulators (e.g., the cytoplasmic tyrosine kinase pp60Src, phosphoinositide 3-kinases) and different receptor tyrosine kinases, such as EGFR, PDGFR, and AXL (7,14). In addition, invadopodia contain various matrix-degrading proteases such as MT1-MMP (MMP14), MMP2, and MMP9 (6,15).

Interestingly, despite the central involvement of invadopodia in the metastatic process and the vast effort invested in their characterization, the molecular mechanisms underlying their formation and activation are still poorly understood. Particularly challenging are the processes whereby cancer cells sense their microenvironment and respond to its chemical and mechanical properties (e.g., its rigidity) by activating the signaling networks that trigger invadopodia formation. Recent studies suggested, somewhat indirectly, an involvement of the Hippo signaling pathway in invadopodia formation. For example, Amotl2 (the angiomotin family member angiomotin-like-2) was shown to localize to podosomes and invadopodia, where it modulates the organization of the actin cytoskeleton (13,16). Interestingly, Amotl2 was also shown to regulate the Hippo signaling pathway by interacting with its transcription co-activator YAP (17). In mammals, core components of this serine/threonine kinase signaling cascade include MST1 and MST2 and the large tumor suppressor kinases (LATS1 and LATS2) that suppress the activity of the transcriptional activators YAP and TAZ (18). YAP/TAZ was shown to act as “mechanosensing switches” that respond to the chemical and physical properties of the cell microenvironment by modulating cellular activity and fate (19,20). Likewise, invadopodia were shown to interact with the ECM via specific integrin receptors (5–8) and to be affected by the rigidity of the underlying matrix (21). Yet, specific information on the involvement of the Hippo pathway in invadopodia formation and activity is limited and mostly indirect. YAP was shown to localize to invadopodia in Src-transformed NIH-3T3 fibroblasts (22), though its involvement in the invasive process remains unclear and lacks mechanistic insights. A direct involvement of YAP through a nucleotide exchange factor TIAM1 was recently described (23)

In this study, we explored the possible roles of YAP and TAZ in invadopodia formation and invadopodia-mediated ECM degradation. Towards this aim, we initially screened a panel of 21 cultured cancer cell lines to check their capacity to form invadopodia in vitro and degrade the underlying gelatin ECM. Suppression of YAP, TAZ or both in most of the invadopodia forming cell lines within this panel, enhanced invadopodia formation and function. A similar effect was obtained when these cells were treated with the YAP/TAZ inhibitor verteporfin. Conversely, overexpression of either YAP, TAZ, or both, effectively blocked gelatin degradation. To identify specific invadopodia-associated components that are affected by YAP and TAZ suppression, we conducted proteomic and transcriptomic profiling of MDA-MB-231 breast cancer cells, that demonstrated the most prominent enhancement of invadopodia following YAP/TAZ depletion. Analysis of the proteomic data revealed that 94 proteins were differentially expressed upon co-knockdown of YAP/TAZ. Among these, nine invadopodia-associated proteins showed significant changes, including an increase in the key core invadopodia components Tks5 and MMP14, which are essential for invadopodia formation (24–27). Furthermore, transcriptome analysis identified 18 differentially expressed invadopodia-related genes following co-knockdown of YAP/TAZ in the same cells. Overall, these results show that YAP and TAZ act as negative regulators of invadopodia formation and matrix degradation in multiple cancer cell lines, suggesting a regulatory role for these transcriptional co-activators in cancer invasion and metastasis.

## Materials and Methods

### Fluorescence Microscopy Reagents

Primary antibodies used in this study included; rabbit monoclonal antibody anti-YAP/TAZ (D24E4; Cell Signaling Technology, Catalog No-8418), rabbit polyclonal anti-TKS5 antibody (Santa Cruz Biotechnology, Catalog No: SC-7390), and rabbit polyclonal anti-TKS5 antibody (Merck, Catalog No: 09-403). Secondary antibodies used here included; peroxidase-conjugated goat anti-mouse IgG (Jackson ImmunoResearch Laboratories, Catalog No: 115-035-003), peroxidase-conjugated goat anti-rabbit IgG (Jackson ImmunoResearch Laboratories, Catalog No: 111-035-144), and goat anti-rabbit IgG Alexa Fluor647 (Thermo Fisher Scientific, Catalog No: A32728). F-actin was stained using Phalloidin-tetramethylrhodamine B isothiocyanate (Sigma Aldrich, Catalog No: P1951). Nuclei were stained using 4’,6-diamidino-2-phenylindole dihydrochloride (DAPI; Sigma Aldrich-Aldrich, Catalog No-D9542).

### Cultured cell lines used in this study

Cancer cell lines purchased from the American Type Culture Collection (ATCC) include MDA-MB-231, NCI-H1299, A375, SKOV-3, OVCAR-3, A549, PC3, PANC-1, MDA-MB-468, A2780, HCC1937, and HCC70. Melanoma cell lines 63T, WM793, CSK-A375, and A2058 were sourced as previously described (28). Cell lines IGR-1, LOX-IMVI, and Malme-3M were kindly provided by Prof. Yardena Samuels (Weizmann Institute of Science, Rehovot, Israel). UM-SCC-47 line was a kind gift from Dr. Itay Tirosh (Weizmann Institute of Science, Rehovot, Israel). Melanoma cell line SB-2 was a kind gift from Prof. Menashe Bar-Eli (The University of Texas MD Anderson Cancer Center, Houston, TX, USA). Cancer cell lines MDA-MB-231, A375, CSK-A375, WM793, A2058, 63T, IGR-1 LOX-IMVI, SB-2, Malme-3M, SKOV-3, OVCAR-3, A549, PC3, PANC-1, MDA-MB-468 and A2780 were cultured in DMEM, supplemented with 10% FCS (Gibco) 2 mM glutamine, 2 mM NEAA, and 100 U/mL penicillin-streptomycin. Cell lines NCI-H1299, HCC1937, and HCC70 were cultured in an RPMI medium containing 10% FCS (Gibco), 2 mM glutamine, 2 mM NEAA and 100 U/mL penicillin-streptomycin. UM-SCC-47 cell line was cultured in three parts of Hams F-12 nutrient mixture medium and one part of DMEM with 10% FCS (Gibco) and 100 U/mL penicillin–streptomycin. All cell lines were maintained at 37°C under a humidified atmosphere of 5% CO_2_ and 95 % air. All cell lines were routinely tested for mycoplasma using the mycoplasma detection kit, MycoAlert™ (Lonza Nottingham, Ltd). Frozen vials of cells stock were thawed, and cells were cultured for no more than a month for the experiments. Additional information on all the cell lines used in this study is summarized in Table S1.

### Transfection procedures

#### Knockdown experiments

Transfection was performed using siGENOME Human YAP1 siRNA, SMART pool; M-012200-00-0005 and siGENOME Human WWTR1 (TAZ) siRNA, SMART pool; M-016083-00-0005 (GE Healthcare Dharmacon). siRNA non-targeting pool #2 (GE Healthcare Dharmacon) was used as a control for all the knockdown experiments. The SMART pool and single oligo siRNA sequences used for the experiments are enlisted in the supplementary Table S2 and Table S3. The final concentration for each siRNA was 30 nmol/L. The siRNAs were transfected using DharmaFect transfection reagent, according to the manufacturer’s protocol. Fresh medium was added after 6 h to replace the medium with transfection reagent and oligonucleotides. Cells were incubated for 48 h following the siRNA transfection and proceeded for further experiments.

#### Overexpression experiments

IRES-GFP empty vector was sourced as previously described (Revach et al., 2019). pcDNA Flag YAP1 and pcDNA3 Flag TAZ were obtained as described previously (29). Flag YAP1 and Flag TAZ were transfected using the jetPEI transfection reagent (Polyplus-Transfection^®^) according to the manufacturer’s instructions. The plasmids were purified using Qiagen Maxi Kit and quantified using a Nanodrop spectrophotometer (Thermo Fisher Scientific, Catalog No: ND-2000). Briefly, MDA-MB-231 and NCI-H1299 (0.7 × 10^6^) cells were seeded on a 10 cm cell culture dish and incubated for 24 h. Subsequently, the cells were transfected with empty vector, Flag-YAP1, Flag-TAZ, and Flag-YAP1 plus Flag-TAZ. The final concentration of each plasmid used for overexpression was 10 μg per 10 cm culture dish. For Flag-YAP1 plus Flag-TAZ condition, the total concentration was 20 μg per 10 cm cell culture dish. The cells were then incubated for 48 h before proceededing with further experiments.

### Preparation of gelatin-coated culture plates

Gelatin coating of culture plates for the matrix degradation assay was conducted as previously described (28). Briefly, ninety-six micro-well plates, with a glass-bottom (Thermo Fisher Scientific Catalog No-164588) were treated with 50 mg/mL of poly-lysine (Sigma Aldrich, Catalog No-P-7405) prepared as 1:1 mixture with Dulbecco’s Phosphate Buffered Saline (DPBS, Biological Industries, Catalog No-02-023-1A) and incubated for 20 minutes at room temperature. Then, the poly-lysine solution was removed, and the plate was washed (×3) with DPBS. Porcine skin gelatin (Sigma Aldrich, catalog no. G2500) was prepared in DPBS (2 mg/ml) and filtered through a 0.22-micron Steritop filter (Millipore Fisher Scientific Catalog No-15770319). Gelatin was fluorescently labeled using Alexa Fluor 488 Protein labeling kit (Molecular Probes, Thermo Fisher Scientific) according to the manufacturer’s instructions. The gelatin was subsequently cross-linked using N-(3-Dimethylaminopropyl)-N’-ethylcarbodiimide hydrochloride (EDC hydrochloride) (Sigma Aldrich Catalog No-03450) and N-Hydroxysuccinimide (NHS) (Sigma Aldrich Catalog No-130672), prepared as 10% solutions in ddH2O. Subsequently, 96-well glass-bottom plates were coated with 40μl volume of a mixture (10:1) of unlabeled gelatin and the Alexa Fluor488 labeled gelatin. For every 100μl gelatin, the gelatin to cross-linker mixture ratios was (82.5μl gelatin: 12.5 μl NHS: 5μl EDC). Subsequently, the gelatin mixture was incubated for 1 h at room temperature. After incubation, the surfaces of the gelatin-coated plates were washed three times with DPBS. 100μl of fresh DPBS solution was added to the wells and UV sterilized for 30 minutes in the biosafety hood. The plates were then ready to use.

### Gelatin degradation assay

The different cancer cell lines (10^4^) were seeded on the Alexa Fluor 488-labeled gelatin matrix in the 96-well plates and cultured for 5-6 h. The cells were then fixed and stained for F-actin and DAPI, washed with DPBS, and kept wet for imaging. Every experiment was performed in two or more replicate wells for each condition. Z stack of images was acquired using a WiScan^®^ Hermes Automated High Content Imaging System (IDEA Bio-Medical Ltd.) using 40X/0.75 NA air objective. In a single well, images from a total of 36 fields were acquired, and the cells were counted based on the nuclear (DAPI) staining. The degraded gelatin area (μm^2^ per cell) was calculated using the Image J software (rsbweb.nih.gov/ij). All degradation values following treatments were compared to those of control cells cultured on the same plate in every independent set of experiments.

### The effect of EGF stimulation on gelatin degradation

Cancer cell lines (SKOV-3 and OVCAR-3) were grown in medium containing 10 % FBS and serum-starved (0.5 % FBS) for 24 h. Subsequently, cells that were cultured in 0.5 % FBS were stimulated with hEGF (recombinant EGF) (Sigma Alrdich, Catalog No: E9644) in concentrations (10 and 30 ng/ml) and plated on Alexa 488-gelatin for 5-6 h. The cells were then processed for the gelatin degradation assay as described above.

### The effect of YAP/TAZ inhibitor verteporfin on gelatin degradation

For gelatin degradation, assay measuring the effect of the YAP/TAZ inhibitor verteporfin (Holland Moran, Catalog No 1711461), MDA-MB-231 and NCI-H1299 cells were seeded on the Alexa Fluor 488 labeled gelatin matrix and cultured for 2 h. Subsequently, the cells were treated either with vehicle (0.1% DMSO) or with different concentrations of verteporfin (final concentration range: 0.5-20 μM) for 4 h. The cells were then fixed and stained for TRITC-Phalloidin and DAPI and imaged using a WiScan^®^ Hermes Automated High Content Imaging System (IDEA Bio-Medical Ltd.) under 10X/0.75 NA air objective.

### Immunostaining, microscopy, and image analysis

Cells (10^4^/well) were seeded on the Alexa Fluor488-tagged gelatin in 96-well glass-bottom plates. At the end of the gelatin degradation experiment (usually 5-6 hours after plating) the cells were fixed and permeabilized for 3 minutes with 3% PFA, 0.5% Triton-X100, in DPBS followed by 3% PFA for another 30 minutes. Then, the cells were washed (×3) with DPBS and incubated with the primary antibody for 1 h. Subsequently, the cells were washed with DPBS (×3) and incubated with the appropriate secondary antibody for 30 minutes. The cells were washed again (×3) with DPBS and kept in DPBS for imaging. The images were acquired using a DeltaVision Elite microscopy system, equipped with a microtiter stage (Applied Precision Inc., Issaquah, WA) with 40X/0.75 air or 60X/1.42 oil objectives (Olympus). All acquired images were analyzed using Image J software (https://imagej.nih.gov/ij/).

### Cell viability assay

A microscopy-based cell viability assay was performed as previously described (30). Hoechst 33342 (1 μg/ml; ImmunoChemistry Technologies, Bloomington, MN, USA) and propidium iodide (250 ng/ml; Sigma Aldrich, St. Louis, MO, USA) in DMEM were added onto cells and kept in the 37 °C incubator for 45 minutes. The cells were centrifuged at 1200 rpm for 3 minutes and then proceeded for imaging. The images were acquired uusing a WIScan Hermes^®^ microscope with a 10X objective (IDEA Bio-Medical Ltd.), and the percentage of live and dead cells was calculated using WiSoft^®^ Athena software (IDEA Bio-Medical Ltd.).

### Western blot analysis

Cells plated on a 10 cm cell culture dish were scraped using a cell scraper and suspended in 300 μl ice-cold RIPA buffer. The cell lysates were kept on ice and vortexed at 5-minute intervals for over 45 minutes. Then, the lysates were cleared by centrifugation at 274 g at 4°C for 10 minutes. The cell lysates were either freshly examined or stored at −80°C. The frozen samples were thawed on ice, subjected to 10% polyacrylamide/SDS gel electrophoresis, and subsequently blotted onto Polyvinylidene fluoride (PVDF) membrane (Merck Millipore^®^ Catalog No IPVH00010). The blots were blocked using 5% skimmed milk in Tris Buffered Saline, containing Tween 20, pH 8.0 (TBST) buffer, and probed with primary antibody overnight at 4 °C. They were then washed with TBST buffer X3 (10 minutes each) and incubated with HRP-coupled secondary antibody (Jackson ImmunoResearch Laboratories Inc.) for 1 h. Chemiluminescent Super Signal West Pico substrate (Thermo Fisher Scientific, Catalog number: 34579) was used for detecting the bands, and the blots were imaged using ChemiDoc MP Imager and quantified using Image Lab 4.1 software (BioRad, USA).

### Quantitative real-time PCR

RNeasy Mini Kit (Catalog No-74104; Qiagen) was used for isolating total RNA from cells. Total RNA (1-2 μg) was reverse-transcribed using LunaScript™ RT SuperMix Kit (New England Biolabs, Catalog No; E3010S). Quantitative real-time RT-PCR (qRT-PCR) was performed using a Fast SYBR Green Master Mix using the OneStep instrument (Applied Biosystems). The obtained values were normalized to either HPRT1 or GAPDH genes. The primers used for the experiments are enlisted in Table S4.

### Preparation of cells for proteomic profiling

For knockdown experiments, MDA-MB-231 (0.7× 10^6^) cells were seeded on 10 cm cell culture dishes, incubated for 24 h, transfected with siControl, siYAP, siTAZ, and a mixture of siYAP and siTAZ SMARTpools, and further incubated for 48 h. The YAP/TAZ knockdown cells were seeded on unlabeled gelatin-coated 10 cm plates and cultured at 37°C for 5-6 h. The effect of the treatment on the cells’ gelatin degradation activity was verified, in parallel, by plating a sample of the same cells on Alexa Fluor 488-gelatin plates and measuring their gelatin degradation phenotype. The experimental design consisted of three independent set of experiments for each knockdown condition. The cells were washed with 5 ml cold PBS and then scraped into 1 ml fresh ice-cold PBS, centrifuged at 3000 rpm at 4 °C, and the cell pellets were flash-frozen in liquid nitrogen and stored at −80 °C.

### Sample preparation for proteomic analysis

Frozen cell pellet samples were dissolved in 5% SDS, 50 mM Tris-HCl (pH 7.5) The total protein concentration was measured using a BCA assay. 100μg of each sample was used for the downstream preparation. Dithiothreitol (DTT) was prepared fresh in 50 mM ammonium bicarbonate, and added to a final concentration of 5 mM. The samples were then incubated at 56°C for 1 hour. Iodoacetamide was prepared fresh in 50mM ammonium bicarbonate, and was added to final concentration of 10mM. Samples were incubated in the dark for 45 min. Phosphoric acid was then added to the samples to final concertation of 1%. The samples were mixed with 350 μL of 90% methanol along with 10% 50 mM ammonium bicarbonate, then transferred to the S-trap filter and centrifuged for 1 min at 4000xg and washed 3 times with 400μL of 90% MeOH+10% 50mM ammonium bicarbonate, then centrifuged at 4000g 1min. 4 μL of 0.5 μg/μL Trypsin in 125 μL in ammonium bicarbonate (25:1 protein amount: trypsin) was added to the samples. Samples were incubated at 37°C overnight. The next day, peptides were eluted using 80μL 50mM ammonium bicarbonate, which was added to the S-trap cartridge, centrifuged at 4000g for 1 min into new tubes and collected the peptides. Then, a second digestion was performed using 4 μL of 0.5 μg/μL trypsin in 50mM ammonium bicarbonate was added to the eluted samples and incubated at 37°C for 4 hrs. Two more elations from the S-trap cartridge were performed. One with 80 μl of 0.2% formic acid, which was added to the S-trap cartridge and spun down at 4000g for 1 min. The second was done using 80 μL of 50%acetonitrile+0.2% formic acid was added to the cartridge and spun down at 4000g for 1 min. The three elutions were mixed and dried using a vacuum centrifuge (Centrivac, LabConco).

### Proteomic analysis

The resulting peptides were analyzed using a **nanoAcquity** liquid chromatography (Waters) coupled with a Q Exactive HF-X (Thermo fisher scientific). Samples were analyzed randomly, loaded on a Symmetry C18 trap column (20mm X 0.18mm, 5um, Waters), and resolved on a HSS T3 (250mm X 0.075mm, 1.8um, Waters) analytical column at 350nl/min, using a gradient of 4-27%B (MeCN, 0.1% formic acid) for 155min. MS1 acquisition was performed at m/z range of 375-1650m/z at 120,000 resolution (@400m/z), allowing Automatic Gain Control (AGC) target of 10^6^ with a maximum Injection Time (IT) of 60ms. MS2 acquisition was performed on the Top10 ions at Data-Dependent Acquisition (DDA) using Higher-energy Collisional Dissociation (HCD) fragmentation set at 27 Normalized Collision Energy (NCE) acquired at 15,000 resolution (@200 m/z). IT was set to 60ms and AGC to 1e5. Dynamic exclusion was set to 30sec with a counter of 1. The resulting data was processed with MaxQuant (v1.6.6.0). The data were searched with the Andromeda search engine against the Human proteome database (SwissProt Nov20) appended with common lab protein contaminants. The following modifications were allowed: fixed carbamidomethylation on C, variable protein N-terminal acetylation, variable deamidation on NQ and variable oxidation on M. The qquantification was based on the LFQ method, based on unique peptides.

### Bioinformatics analysis

For each cancer cell lines used in this study (See Supplementary Table S1), we retrieved the information concerning the tissue of origin (e.g., primary tumor vs. metastases) from the CCLE database (https://depmap.org/portal/download/). Bioinformatic analysis of the proteomic data of MDA-MB-231 cells was applied on LFQ intensities of 4,704 detected proteins. Proteins having at least two one razor and unique peptides were considered, and 37 known contamination were removed from the analysis. For the detection of differential proteins, intensities were log2 transformed and analyzed with ANOVA following a multiple test correction (FDR step-up) using Partek Genomics Suite 7.0. For each pairwise comparison, we considered proteins having at least two valid measurements (out of 3) in both groups and that passed the thresholds of fold change (log2)| > 1 and p-value <0.16. In addition, proteins that were detected in at least 2 replicates in one group and completely absent in the other group were also considered as qualitatively differential proteins. For visualization of protein expression, heat maps were prepared using Partek Genomics Suite, using log2-transformed LFQ intensities with row standardization (scaling the means of a row to zero, with a standard deviation of 1), and partition clustering using the k-means algorithm (Euclidian method). For volcano plot visualization, missing values were imputed to a value of 15, and new fold change values were calculated with ANOVA and visualized using MATLAB. Principle component analysis (PCA) was calculated using Partek Genomics Suite. For visualization of proteins network, the relations between differential proteins of the double knockdown was inferred with StringDB (31) as a “full STRING network”, and visualized by Cytoscape 3.7.2 (32). The width of the edges corresponds to the “combined score” or StringDB, and the protein color scale corresponds to their log2 fold change as inferred from the proteomics ANOVA. Proteins that change qualitatively (were detected only in one condition) were assigned an imputed value of +/− 5. The assignment of proteins belonging to Hippo signaling, cell adhesion, and ECM remodeling pathways was inferred using the GeneCards suite (33). A list of invadopodia-related proteins was compiled by data mining in the Harmonizome database and the related literature (34–43).

### RNA sequencing

MDA-MB-231 cells (0.7 ×10^6^) were seeded on 10 cm culture dishes and incubated for 24 h. Subsequently, the cells were transfected with SMARTpool siRNA for siControl, siYAP, siTAZ, and siYAPTAZ and incubated for a further 48 h for the knockdown. After the incubation, the knockdown cells were seeded on non-labeled gelatin-coated 10 cm plates and cultured for 5-6 h in an incubator. Subsequently, the culture medium was aspirated, and the cells were given a wash with DPBS. One ml of DPBS was added to the plate, and the cells were then scraped using a cell scraper. Then the cells were centrifuged at 3000 rpm at 4 °C. The supernatant was aspirated, and the RNA was extracted from the cell pellet. The RNA extraction was done using RNeasy Mini Kit (Catalog Nos; 74104, 74106 Qiagen) according to the manufacturer’s instructions. The RNA was quantified using Qubit 3 Fluorometer (ThermoFisher scientific, USA) and TapeStation (Agilent Technologies 4200, USA) to assess the purity of RNA. A RIN (RNA Integrity Number) score ranging from 9-10 was obtained for each condition. Then RNA-seq libraries were prepared at the Crown Genomics institute of the Nancy and Stephen Grand Israel National Center for Personalized Medicine, Weizmann Institute of Science. Libraries were prepared using the INCPM-mRNA-seq protocol. Briefly, the polyA fraction (mRNA) was purified from 500 ng of total input RNA followed by fragmentation and the generation of double-stranded cDNA. Afterwards, Agencourt Ampure XP beads cleanup (Beckman Coulter), end repair, A base addition, adapter ligation and PCR amplification steps were performed. Libraries were quantified by Qubit (Thermo fisher scientific) and TapeStation (Agilent). Sequencing was done on a NovaSeq6000 instrument (Illumina) using an SP 100 cycles kit (single read sequencing).

### Transcriptomic analysis

RNA sequencing analysis was done using the UTAP transcriptome analysis user-friendly Transcriptome Analysis Pipeline (UTAP v1.10) transcriptome analysis pipeline (Kohen, Barlev et al. 2019). Reads were trimmed to remove adapters and low quality bases using cut adapt (-a “A pipeline (44). Reads were trimmed to remove adapters and low quality bases using cut adapt (-a “A (10)” -a “T (10)” –times 2 -q 20 -m 25) (45) and mapped to the human genome (GRCh38, GENECODE version 34) using STAR v2.4.2a (46) (using–alignEndsType EndToEnd, –outFilterMismatchNoverLmax. 0.05, –two pass Mode Basic). Reads were counted using STAR, and genes having minimum of five reads in at least one sample were considered. Normalized counts and detection of differential expression were performed using DESeq2 (47) (betaPrior, cooksCutoff, and independent filtering parameters set to False). Differentially expressed genes were selected with absolute fold change (log2)≥1 and adjusted multiple testing p-value ≤0.05 (48). Matlab was used to generate the volcano plots.

### Data Availability

The raw data of proteomic profiling has been deposited at the ProteomeXchange via the Proteomic Identification Database (PRIDE partner repository). The RNA-seq data has been deposited in in NCBI’s Gene Expression Omnibus (Edgar et al., 2002) and are accessible through Genome Sequence Archive for Human. There are no restrictions on the availability of the data.

### Statistical analysis

All statistical analysis of the experimental data was performed using the Graph Pad Prism version 8.0.1 for Windows, GraphPad Software, San Diego, California USA, www.graphpad.com software. Statistical significance for each experiment is marked in the form of asterisks (*) along with the calculated p-value for experiments are shown.

## Results

### Differential gelatin degradation and invadopodia formation by a panel of cultured cancer cell lines

Towards the selection of cultured cancer cell lines that would be suitable for testing the involvement of the Hippo pathway in invadopodia formation and matrix degradation, we assembled a panel of 21 cancer cell lines (See Supplementary Table S1). These lines were derived from the following cancers: melanomas (A375, CSK-A375, A2058, WM793, 63-T, IGR-1, Malme-3M, SB-2, LOX-IMVI); breast carcinoma (MDA-MB-231, MDA-MB-468, HCC1937, HCC70); head and neck squamous cell carcinoma (UM-SCC-47); non-small cell lung carcinoma (NCI-H1299); lung adenocarcinoma (A549); ovarian carcinoma (SKOV-3, OVCAR-3, A2780); prostate carcinoma (PC-3) and pancreatic cancer (PANC-1). For details on the tissue of origin, site of isolation (primary tumor vs. metastasis) and capacity to degrade gelatin matrix, see Supplementary Table S1. Imaging of the gelatin degradation by the 21 tested cell lines revealed 12 cell lines that display a conspicuous gelatin degradation pattern (Figure 1) with an average gelatin degradation score >1 μm^2^/cell. Quantification of the gelatin degradation by these cell lines pointed to high variability, ranging from an average degradation score of 3 μm^2^/cell to over 100 μm^2^/cell (Figure 2). It is noteworthy that the 12 “degradation-positive” cell lines were originally derived from melanoma (9/9), breast cancer (1/4), NSCLC (1/1), and squamous cell carcinoma (1/1). Furthermore, the gelatin degrading cell lines formed conspicuous actin-rich invadopodia, often overlapping with the degraded (dark spots) areas on the gelatin matrix (Figure 1 and “zoomed-in” images of IGR-1 and SB-2 cells, shown in Supplementary Figure S1). The nine cell lines that displayed low gelatin degradation (< 1 μm^2^/cell) included those derived from ovarian cancer (3/3), breast cancer (3/4), lung adenocarcinoma (1/1), pancreatic carcinoma (1/1), and prostate carcinoma (1/1), (Supplementary Figure S2). Treatment of two of these nine cancer cell lines (SKOV-3 and OVCAR-3) with EGF (10 and 30 ng/mL, see also (49) did not trigger enhanced invadopodia formation or gelatin degradation in these cells (Supplementary Figure S3).

**Figure 1.**
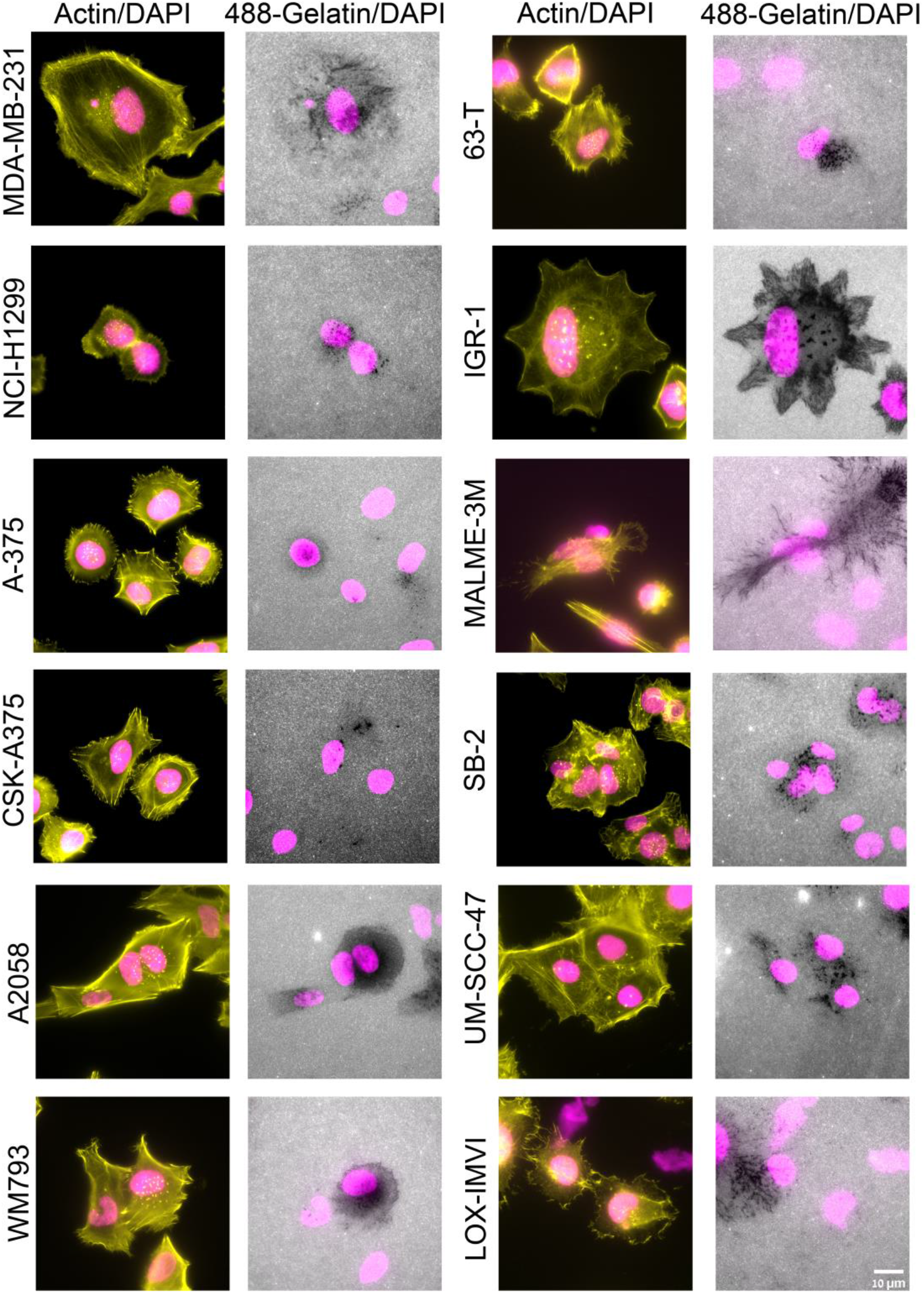
A panel of multi-cancer cell lines screened for gelatin degradation and actin rich invadopodia structures. Cell lines were cultured on a fluorescently labeled (488) gelatin and cultured for 5-6 h. Subsequently, the cells were fixed and stained for actin (TRITC-phalloidin) and DAPI. Images were acquired under 60 x/1.42 oil objective. Among the 21 multi-cancer cell line panel used in the study, 12-cancer cell lines (shown here) were found to be positive for the actin rich invadopodia appearing as prominent actin-rich dots near the nucleus and gelatin matrix degradation. Scale bar is 10 μm.

**Figure 2.**
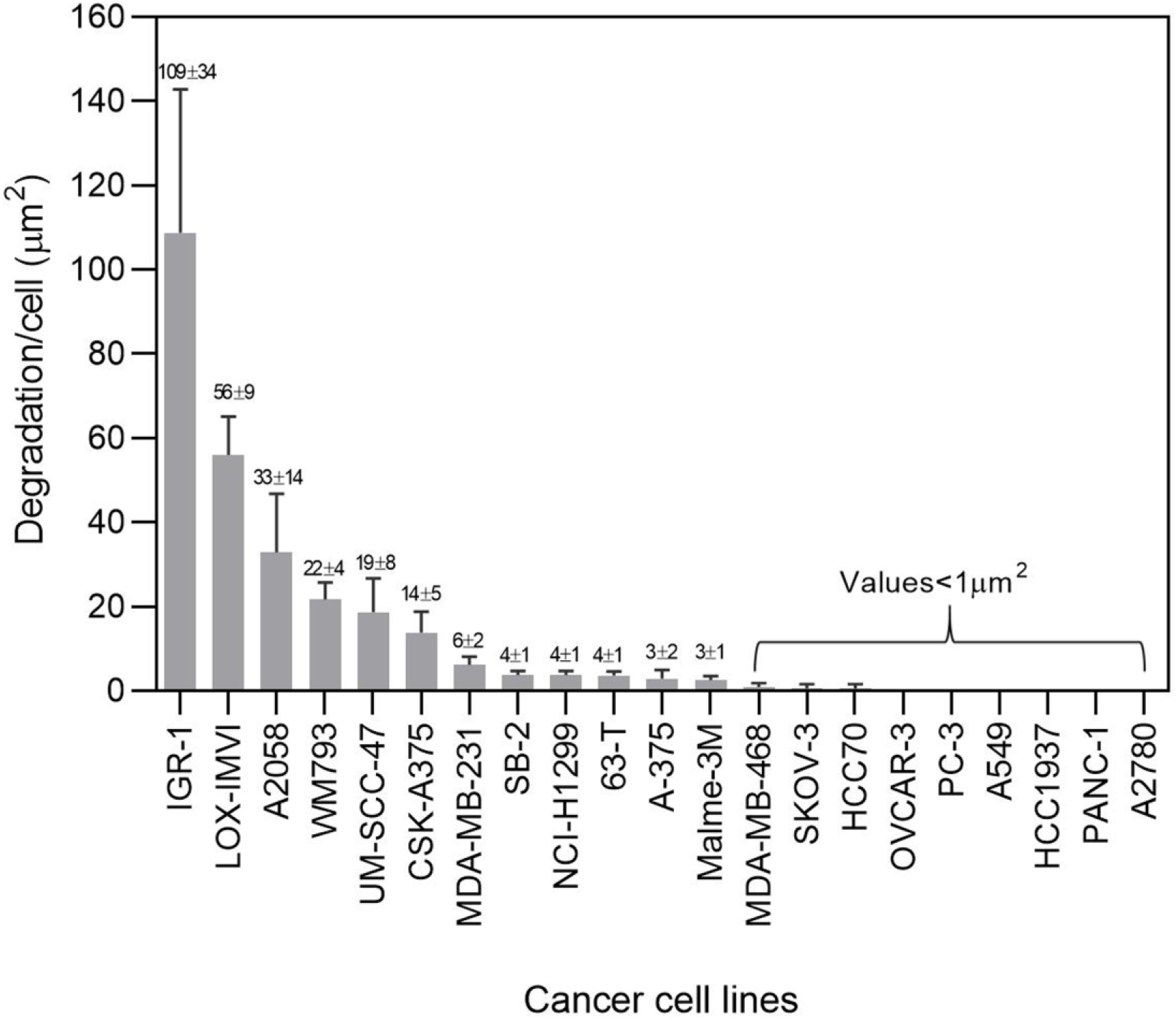
Quantitation of gelatin matrix degradation in a multi-cancer cell line panel. Cells were seeded on FITC labelled gelatin matrix and cultured for 5-6 h. Subsequently, the cells were fixed with 3 % paraformaldehyde in PBS and stained for actin and DAPI. Each cell line was seeded in duplicate wells. Images were acquired using 40x/0.75 air objective. 36 fields were imaged per well and the gelatin degradation/cell (μm^2^) was quantitated using ImageJ software. The plot shows the average of four independent data sets with ± SEM. The p-value was found to be (<0.0001). Four independent sets of experiments were performed and the average plot is shown. These cell lines then served as platform for further experiments in this study.

### Depletion or inhibition of YAP and TAZ enhance invadopodia formation and gelatin degradation in multiple cancer cell lines

To investigate the involvement of the Hippo pathway in invadopodia formation and function, we subjected the entire panel of the 21 cancer cell lines (both those scored positive and those scored negative for matrix degradation and invadopodia formation) to siRNA-mediated knockdown of YAP, TAZ or both (SMARTpool siRNAs are listed in Table S2). As shown in Supplementary Figure S4, the 9 cell lines that displayed average gelatin degradation levels below 1μm^2^/cell remained negative also after YAP and TAZ knockdown. Efficient knockdown was confirmed at both mRNA (Supplementary Figure S6A) and protein (Supplementary Figure S6B) levels, as shown for MDA-MB-231 cells. Off-target effects were excluded by the use of individual siRNA duplexes (sequences in Supplementary Table S3). The effect on the gelatin degradation score is shown in Supplementary Figure S5. Quantification of these results pointed to a significant increase in matrix degradation following knockdown of both YAP and TAZ (about 4-fold). Suppression of either YAP or TAZ yielded more modest, yet consistent elevation of matrix degradation. Similar trends, albeit with varying intensities, were observed also in the other cancer cell lines (Figure 3G). Strikingly, in MDA-MD-231 cells, depletion of YAP or TAZ led to substantial increase in gelatin degradation in most of the cancer cell lines tested, as shown in Figure. 3A and quantified in Fig. 3C.

**Figure 3.**
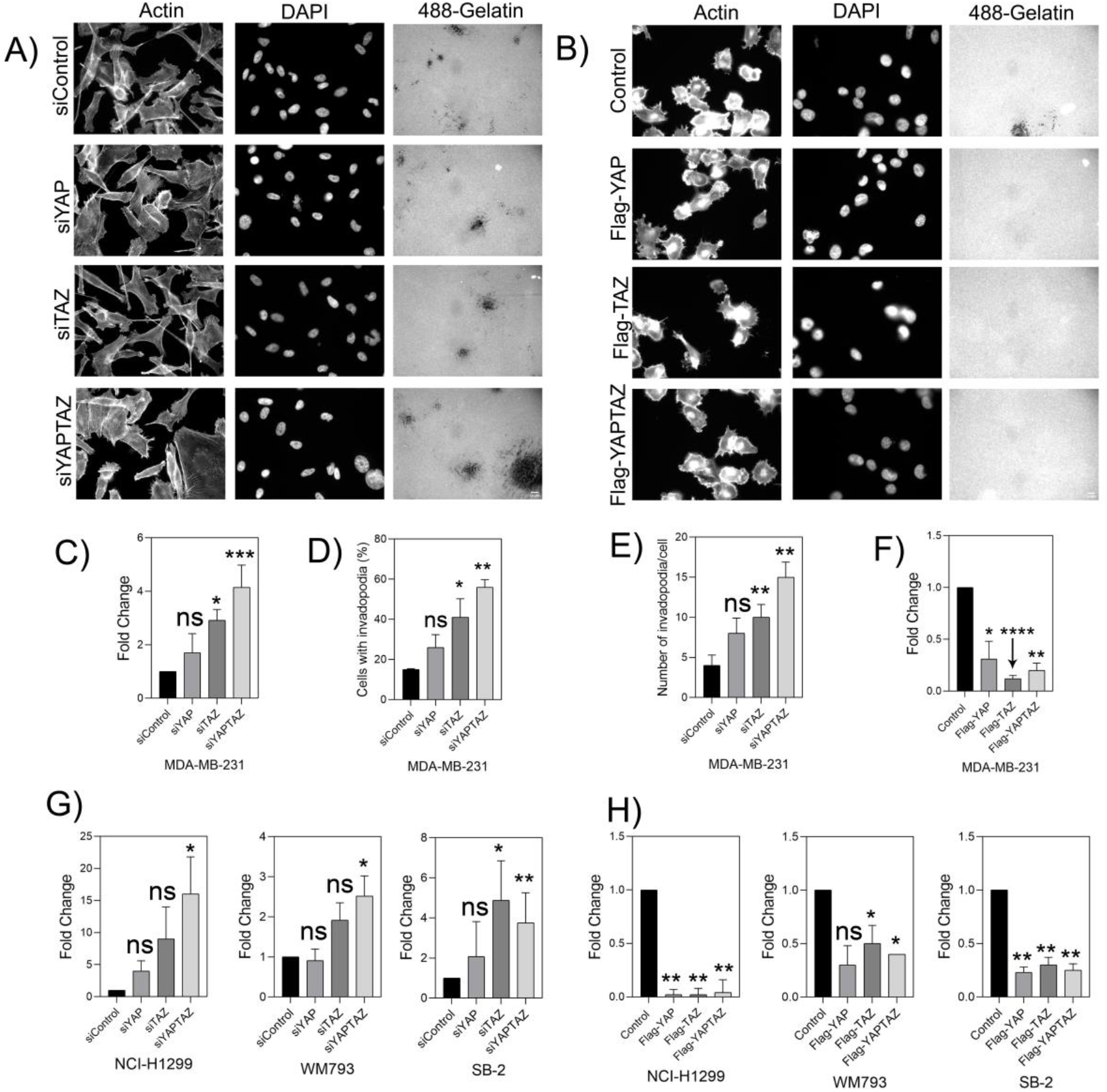
Knockdown of YAP/TAZ elevates gelatin matrix degradation and invadopodia formation in MDA-MB-231. A). YAPTAZ was knockdown for 48 h using SMART pool siRNA. The knockdown cells were then seeded on Alexa Fluor 488-labeled gelatin matrix and cultured for 5-6 h. Cells were fixed and stained for actin and DAPI. Z-stack of images for actin, DAPI and Alexa Fluor 488-gelatin were captured under 40 X (0.75) air objective using a WIScan Hermes^®^ microscope (IDEA Bio-Medical Ltd.). Four independent set of experiments were performed. A representative panel of the captured images are shown. Scale bar is 10 μm. B). Cells were overexpressed with empty vector control, Flag-tagged YAP, TAZ and YAP+TAZ plasmids and incubated on Alexa Fluor 488-gelatin for 5-6 h. Cells were fixed and stained for actin and DAPI. Z-stack of images for actin, DAPI and Alexa Fluor 488-gelatin were captured under 40 X (0.75) air objective using a WIScan Hermes^®^ microscope (IDEA Bio-Medical Ltd.). Four independent set of experiments were performed. A representative panel for the images of one of the set is shown. Scale bar is 10 μm. C). Fold change obtained from the values for gelatin degradation/cell (μm^2^) is plotted for each treatment condition. The significance values calculated by paired t test was (Control vs siYAP) ns, not significant, (Control vs siTAZ) *, *P* value <0.00330, (Control vs siYAP+TAZ) ***, *P* value <0.0008. The *P* value was **** <0.0001 by one-way ANOVA for treatment conditions when compared to control. D). An average plot (Mean ± SEM) of percentage of cells with invadopodia counted for each treatment condition is shown. The significance values for the treatment conditions as calculated by paired t test was (Control vs siYAP) ns, not significant, (Control vs siTAZ)*, *P* value <0.0039, (Control vs siYAP+TAZ)**, *P* value <0.0021. *P* value was *, 0.0251 for treatment conditions when compared to control as determined by one-way ANOVA. E). An average plot (Mean± SEM) of number of invadopodia/cell counted for each treatment condition. The significance values for the treatment conditions as calculated by paired t test was (Control vs siYAP) ns, not significant, (Control vs siTAZ) **, *P* value <0.0032, (Control vs siYAP+TAZ) **, *P* value <0.0020. *P* value was *, 0.0116 for treatment conditions when compared to control as determined by one-way ANOVA F). An average plot of gelatin degradation/cell (μm^2^) with (Mean± SEM) for each treatment condition after overexpression of YAP, TAZ or both is shown. The significance values for the treatment conditions as calculated by paired t test was (Control vs siYAP) *, 0.0264, (Control vs siTAZ)****, *P* value <0.0001, (Control vs siYAP+TAZ)**, *P* value <0.0019. *P* value was *, 0.0197 for treatment conditions when compared to control as determined by one-way ANOVA. G). Fold change obtained from the values for gelatin degradation/cell (μm^2^) for different cancer cell lines after knockdown of YAP, TAZ or both are shown. Data is average (Mean ± SEM) of three independent sets of experiments. (NCI-H1299-ns, not significant, *, *P* value 0.0146 for treatment conditions when compared to control as determined by one-way ANOVA), WM793-*, *P* value 0.0040 for treatment conditions when compared to control as determined by one-way ANOVA, SB-2-* *P* value 0.0197, **, 0.0052 for treatment conditions when compared to control as determined by one-way ANOVA) H). Fold change obtained from the values for gelatin degradation/cell (μm^2^) for different cancer cell lines after overexpression with YAP, TAZ or both are shown. Data is average (Mean± SEM) of four independent sets of experiments. (NCI-H1299-**, *P* value 0.0082 for treatment conditions when compared to control as determined by one-way ANOVA), WM793-*, *P* value 0.00373 for treatment conditions when compared to control as determined by one-way ANOVA, SB-2-**, *P* value 0.0041 for treatment conditions when compared to control as determined by one-way ANOVA) after overexpression with YAP, TAZ or both are shown.

To assess directly the effect of YAP/TAZ knockdown on invadopodia formation we quantified the prominence of invadopodia-containing cells (Figure 3D) and the average number of invadopodia detected in the invadopodia-forming cells (Figure 3E). The quantification of actin-rich invadopodia was based on supervised segmentation of phalloidin-labeled cells (see Materials and Methods). To further confirm that the segmented actin-rich dots are indeed *bona fide* invadopodia, we labeled the cells for Tks5, a highly specific invadopodia component. Comparison of the actin labeling and the Tks5 labeling confirmed that the actin-rich structures are indeed invadopodia (Figure 4). As a complementary approach to YAP/TAZ downregulation, we treated the cells with verteporfin, a potent inhibitor of YAP and TAZ (50,51). Exposure of MDA-MB-231 cells for 6 h to increasing concentrations of verteporfin (0.5-20 μM) had no apparent effect on cell viability. However, treatment with 10 and 20-μM verteporfin resulted in a 3- and 9-fold elevation, respectively, in the gelatin degradation activity (Figure 5). This further supports the notion that YAP and TAZ suppress invadopodia formation and matrix degradation. As shown in Supplementary Figure S4, the 9 cell lines that displayed average gelatin degradation levels below 1μm^2^/cell remained negative also after YAP and TAZ knockdown. Together these experiments show that both YAP and TAZ have a capacity to suppress invadopodia-mediated matrix degradation.

**Figure 4.**
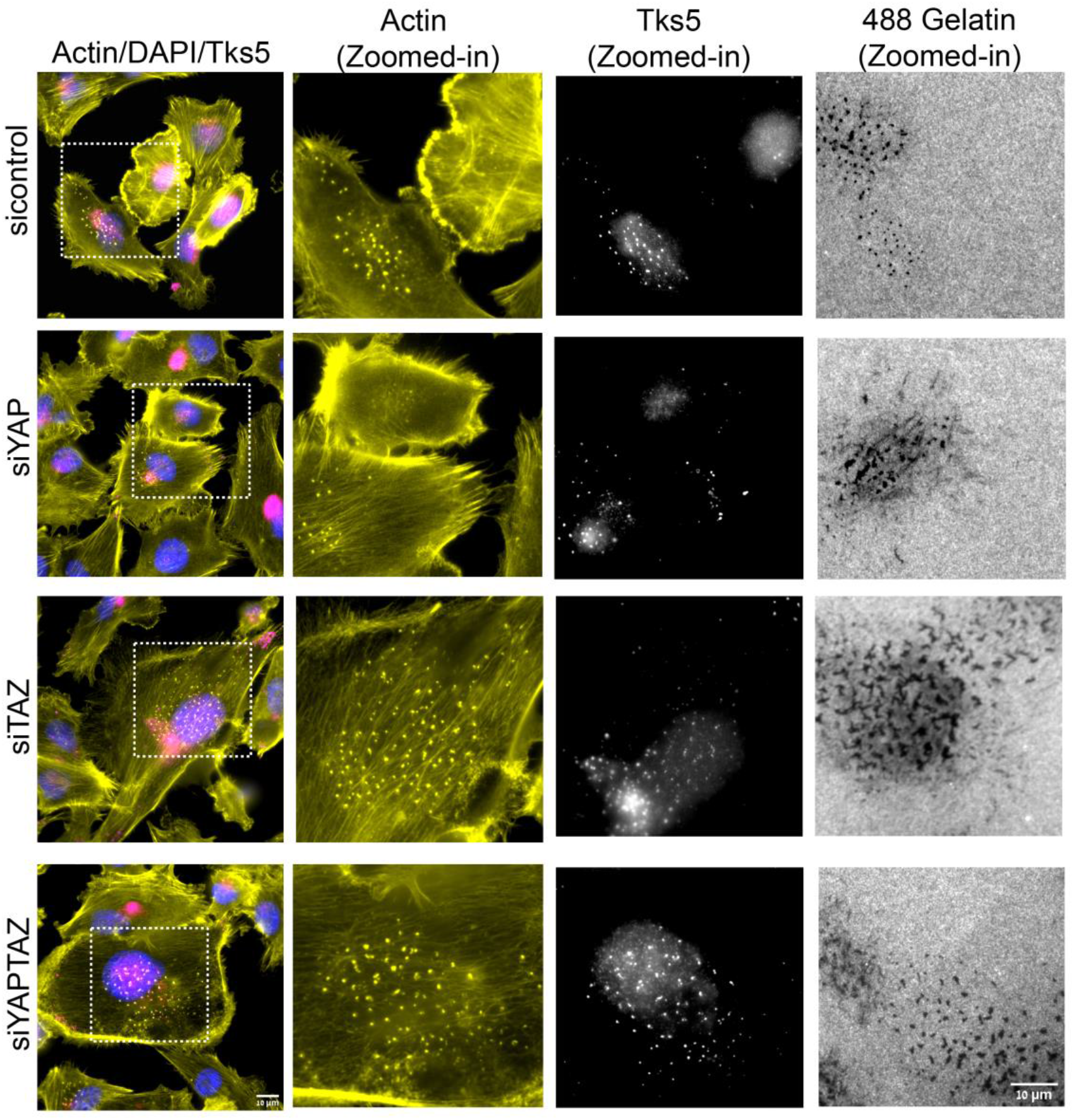
Knockdown of YAP, TAZ or YAP+TAZ in MDA-MB-231 causes elevation in gelatin degradation and invadopodia. MDA-MB-231 cell lines were knockdown for siControl, siYAP, siTAZ and siYAP+TAZ and incubated on Alexa Fluor 488-gletain for 5-6 h. Cells were fixed and stained for actin, DAPI, and invadopodia specific marker Tks5. Cells in the first panel are marked with dotted line white boxes displaying actin (yellow), Tks5 dots (white) and corresponding dark gelatin degradation areas as zoomed in images are shown.

**Figure 5.**
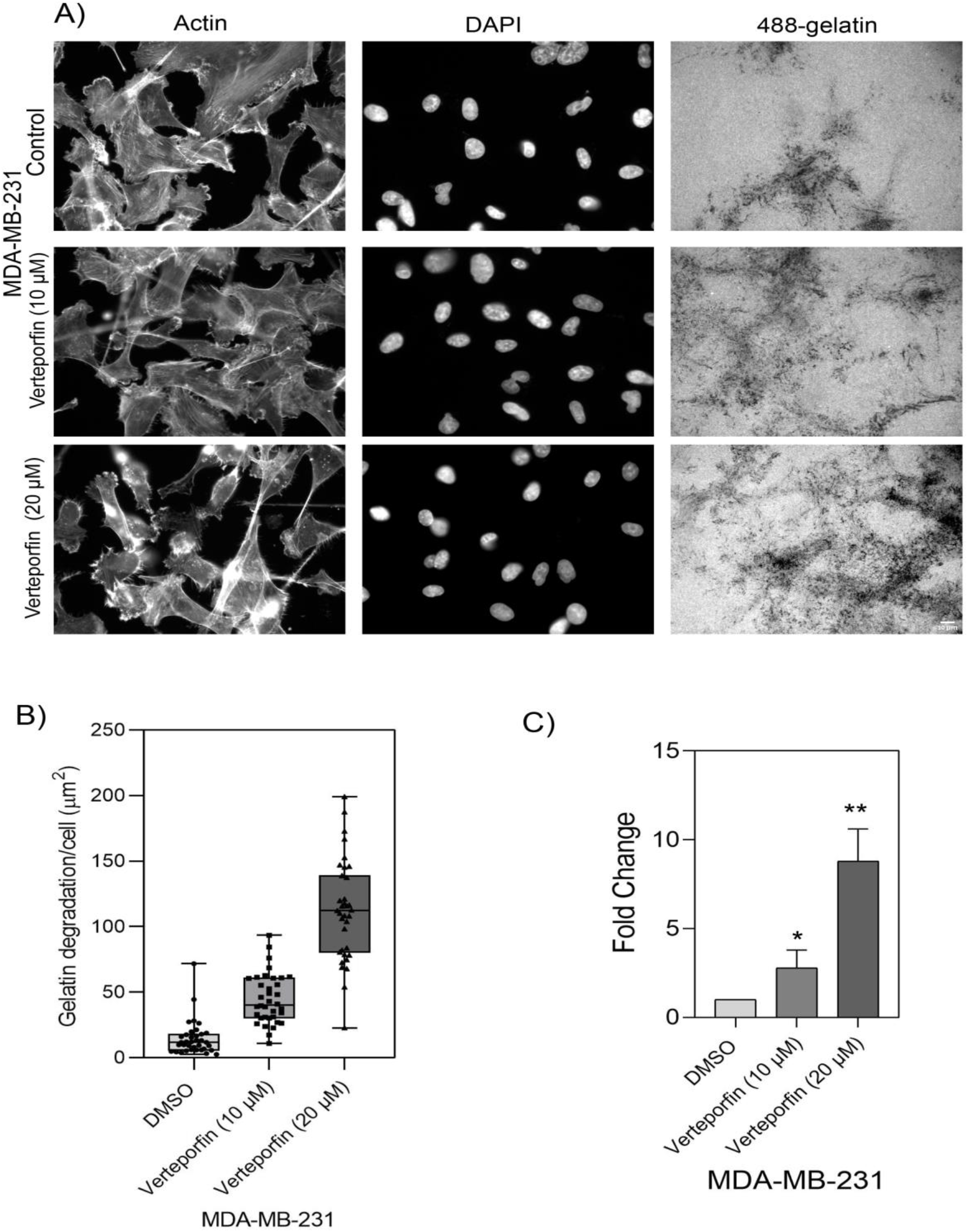
Verteporfin inhibitor causes elevation in gelatin degradation in MDA-MB-231 cell line. MDA-MB-231 cell line was treated with vehicle control, 10 and 20 μM of verteporfin and incubated for 5-6 h on Alexa 488-labeled gelatin. Z-stack of images for actin, DAPI and Alexa Fluor 488-gelatin were captured under 40 X (0.75) air objective using a WIScan Hermes^®^ microscope (IDEA Bio-Medical Ltd.). Four independent set of experiments were performed. A). A representative panel of images for each treatment condition are shown. Scale bar is 10 μm. B). An average plot of gelatin degradation/cell (μm^2^) with ± SEM for each treatment condition is shown. C). Fold change obtained from the values for gelatin degradation/cell (μm^2^) is plotted for each treatment condition. The significance values for the treatment conditions was calculated by two-tailed t test. **, *P* value <0.0033, *, *P* value <0.0408. P value <0.0012 (**) as determined by one-way ANOVA.

### Overexpression of YAP and TAZ suppresses extracellular matrix degradation

To further substantiate the ability of YAP and TAZ to suppress invadopodia-mediated matrix degradation, we overexpressed Flag-tagged YAP, TAZ or both in four “responding” cell lines (MDA-MB-231, H1299, WM793 and SB2), and a Western blot is shown in Figure S6C. As shown in Figure 3B and quantified in Figure 3F and 3H for (MDA-MB-231), (NCI-H1299), (WM793) and (SB-2), overexpression of YAP, TAZ and particularly both together, led to substantial reduction in gelatin degradation as compared to the empty vector control. Overall, our results imply that YAP and TAZ inhibit invadopodia formation in multiple cancer cell lines.

### Proteomic profiling of YAP/TAZ knockdown in MDA-MB-231 cells

To identify downstream molecular targets underlying the apparent suppressive effect of YAP and TAZ on invadopodia formation and degradation we conducted proteomic profiling in MDA-MB-231 cells, which showed the most prominent results. Specifically, the samples subjected to the proteomic analysis consisted of three independent sets of experiments involving: (1) untransfected control cells; (2) siControl (“transfection control”), (3) siYAP, (4) siTAZ, (5) siYAP+siTAZ. This proteomic profiling yielded quantitative data for 4667 proteins (see Supplementary Table S5). Principal component analysis (PCA) of the data revealed that, overall, the control groups (siControl) could be well separated (with PC1) from the sample groups (siYAP, siTAZ, siYAP+TAZ), and that the double knockdown was clearly separated from the single knockdown replicates (Supplementary Figure S7A). Differences in protein expression between the experimental groups were evaluated with ANOVA, using thresholds as described in Materials and Methods. Comparison of the analyses of the “non-transfected” and siControl-transfected cells resulted in only a few (16) differential proteins between the two control groups, suggesting that the transfection might have a small, negligible effect. Thus, further analyses of the knockdown effects were conducted by comparing the siControl and the “Hippo-suppressed” groups (siYAP, siTAZ, siYAP+siTAZ). Altogether, the proteomic profiling analysis detected 122 proteins that were differentially affected upon silencing of either YAP (29 proteins) or TAZ (27 proteins), and a more pronounced effect was detected upon co-knockdown of YAP+TAZ (94 proteins, Supplementary Table S6,). A Venn diagram showing the overlap of differentially expressed proteins in each condition is shown in Supplementary Figure S9. Among the differentially expressed proteins in the double knockdown were proteins known to be involved in the Hippo signaling pathway such as TGFB2, AJUBA, FRMD6, YAP1, SERPINE1, and proteins associated with cell adhesion and ECM remodeling pathways (e.g. SERPINE1, THBS1, COL6A1, MMP14, TIMP3, LAMA5, ITGB5). These results support the role of the Hippo pathway in molecularly regulating ECM degradation.

### YAP and TAZ modulate the levels of key invadopodia-associated proteins

To determine whether knockdown of YAP+TAZ affects the levels of invadopodia associated proteins, we compiled a list of proteins associated with invadopodia structure and functionality (Supplementary Table S7). The list was primarily based on the scientific literature (35,40–42), the *Harmonizome* database (34) and selected proteomic reports (36–39,43). Given that these resources are based on data obtained by different methods, diverse cell types, and that association with invadopodia (e.g., structural, functional, and regulatory) was not strictly and uniformly defined, our list is rather comprehensive albeit with limited editing. This crude list comprises diverse proteins, including membrane receptors, cytoskeletal proteins, adaptor proteins, proteinases and other enzymes, along with the components of different signaling networks. Notably, co-knockdown of YAP and TAZ in MDA-MB-231 cell line revealed significant differential expression of nine invadopodia-associated proteins. Of those, seven proteins (ADAM19, DIAPH2, GSN IDH1, ITGB5, MMP14 (MT1-MMP) and SH3PXD2A (commonly referred to as Tks5) were upregulated whereas two proteins (SERPINE1 and AKAP12) were downregulated. The majority of these proteins are known to play key roles in invadopodia formation, matrix degradation, matrix adhesion and organization of the actin cytoskeleton (Figures 7, 8 and Supplementary Table S8). To determine whether the changes in the invadopodia-associated proteome, described here, were regulated at the transcriptional or the post-transcriptional levels, we conducted transcriptome profiling after co-knockdown of YAP and TAZ in MDA-MB-231 cells (Supplementary Table 9). PCA analysis of the RNA sequencing results showed good separation between the siControl group and the treated groups (siYAP, siTAZ, siYAP+TAZ) (Figure S6B). Similar to the protein profiling results, a more pronounced differential effect of YAP+TAZ knockdown on gene expression was observed, affecting 594 genes. The majority of these genes were affected only when the expression of both YAP and TAZ was suppressed, whereas single knockdown of YAP affected 34 genes and that of TAZ 61 genes (Figure 6 B and Figure S4B). Out of these YAP+TAZ-affected genes, 18 had a documented association with invadopodia, including 5 integrin chains, matrix metalloproteinases, matrix components, signaling molecules and cytoskeletal regulators. Notably, only gelsolin appeared in both the proteomic and the transcriptomic lists of differentially affected invadopodia-related proteins (Supplementary Table 10). Hence, YAP and TAZ are apparently affecting invadopodia formation via multiple regulatory mechanisms.

**Figure 6.**
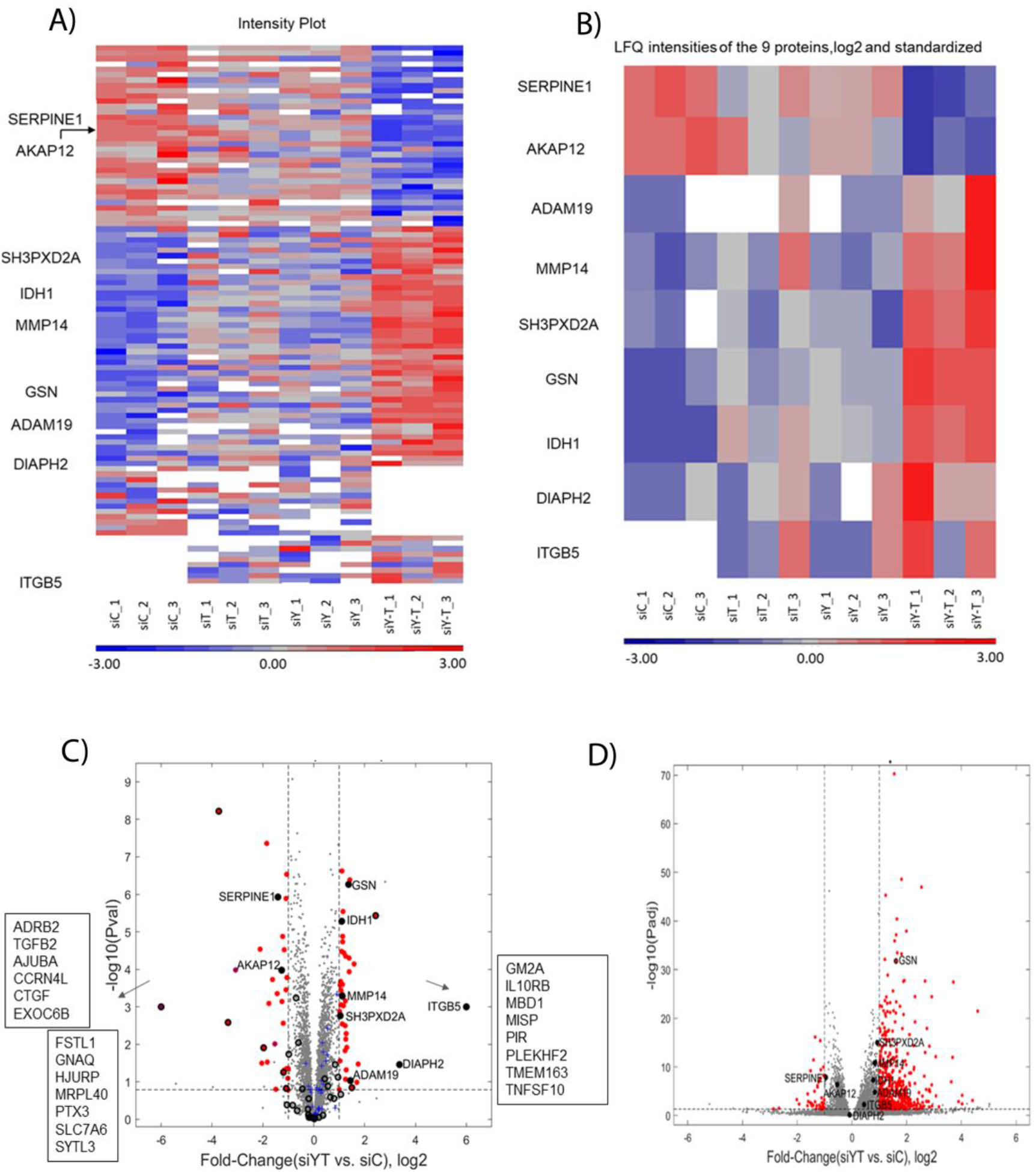
Protein profiling results after knockdown of YAP, TAZ or both in MDA-MB-21 cell line. A). A heatmap of the expression levels of 122 differential proteins obtained from the proteomic profiling for each treatment condition with three independent sets of experiments are shown. LFQ intensities were log2 transformed and standardized. Missing values are shown in white. B) A heatmap of the nine invadopodia-related proteins that were differentially expressed proteins after co-knockdown of YAP+TAZ C). Volcano plot from protein profiling results showing significantly differential proteins marked as red dots indicating up-regulation of proteins such as ADAM19, DIAPH2, GSN, IDH1, ITGB5, MMP14 (MT1-MMP), SH3PXD2A on the right side whereas downregulation of proteins AKAP12 and SERPINE1 on the left side. Proteins that changed qualitatively in the analysis are listed in boxes Blue crosses indicate hippo-signaling pathway proteins detected from the protein profiling results. Black empty circles denote proteins that were detected in the single knockdown of either YAP or TAZ D). Volcano plot for the genes obtained from the transcriptome analysis. Red dots indicate differentially expressed genes.

**Figure 7.**
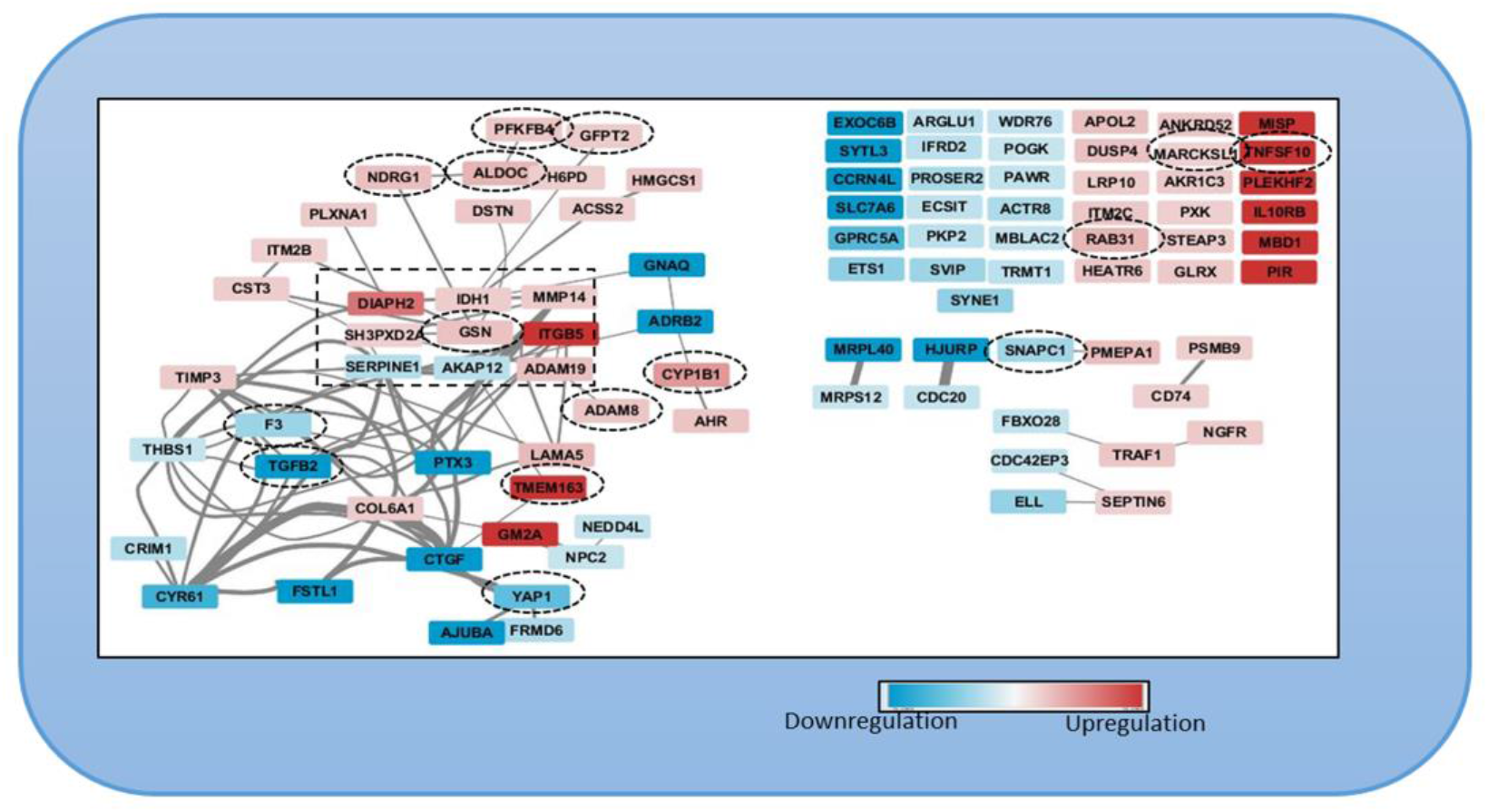
Network of 94 invadopodia specific proteins resulting from the protein profiling analysis after co-knockdown of YAP+TAZ in MDAMB-231 cell line. Invadopodia specific proteins that were up-regulated (red) and downregulated proteins (blue) are shown. 63 proteins had either functional or physical protein associations whereas 38 proteins did not have a known protein association. Proteins differentially expressed in both proteomics and transcriptomic profiling are encircled in black dotted boxes.

## Discussion

The involvement of Hippo signaling in cancer development, progression, and metastasis, has attracted considerable interest in recent years (19,52,53), yet the nature of its effects on the multiple manifestations of the malignant phenotype remains controversial (Ou, Sun et al. 2017, Jho 2018). In the present study, we chose to focus on the mechanisms regulating cancer invasion and metastasis, mediated by matrix-degrading invadopodia. Specifically, we tested the effects of YAP/TAZ suppression or over-expression on the formation of invadopodia and the consequent matrix degradation. To select appropriate cells for these experiments, we have conducted a survey of 21 cultured cancer cell lines derived from diverse primary tumors (10) and metastatic lesions (11). To our knowledge, this is the first and largest comparative characterization of matrix degradation competence of multiple cancer cell lines. Among which, 12 displayed consistent gelatin degradation capacity, with a broad range of degradation scores (~ 3-109 μm^2^/cell) and 9 cell lines that failed to degrade the underlying gelatin. Morphological examination of the gelatin degrading cells revealed actin-rich invadopodia, whereas cells that failed to degrade the matrix were largely devoid of invadopodia (Figures S2). Multi-color fluorescence microscopy further validated that the structures identified as invadopodia, indeed contain TKS5, an invadopodia-specific adaptor protein that often co-localizes with gelatin-degradation sites (Figure 4 and Supplementary Figure 1). Notably, the co-localization of invadopodia with locally degraded matrix is partial, due the dynamic nature of invadopodia formation and turnover. Interestingly, essentially all the cell lines, derived from melanoma cancers (9/9, derived either from metastases or primary tumors) formed conspicuous invadopodia, while cell lines derived, for example, from ovarian adenocarcinomas or breast adenocarcinoma were mostly invadopodia negative (4/4 and 3/4, respectively). Further attempts to induce invadopodia formation and matrix degradation in the invadopodia-negative cell lines, e.g. by changing the environmental conditions (e.g. EGF stimulation; see (49)), or the cells’ signaling machinery (e.g. knockdown of YAP and TAZ, inspired by the present study) failed to turn-on invadopodia formation in these cells, suggesting that these cells might use invadopodia-independent mechanisms for their invasion and dissemination, at least under culture conditions. Suppressing the expression of YAP, TAZ or both in the invadopodia-forming cells, had variable effect on matrix degradation; cells displaying either high or modest gelatin degradation score (e.g. A2058, UM-SCC-47, LOX-IMVI, A375, CSK-A375 and 63-T), were not affected by YAP/TAZ or were partially suppressed, suggesting a Hippo-insensitive regulation of invadopodia in these cells. In contrast, a group of 4 cell lines (MDA-MB-231, H1299, WM-793 and SB2) displayed a pronounced and consistent elevation in invadopodia formation and gelatin degradation following YAP/TAZ knockdown (Figure 3), suggesting that the Hippo pathway (via the inhibition of YAP and TAZ by MST/LATS) promotes invadopodia formation. This view is further corroborated by the enhancement of invadopodia after treating the cells with the YAP/TAZ signaling inhibitor verteporfin (Figure 5) and by the suppression of invadopodia formation and activity following YAP/TAZ over-expression. Interestingly, contrary to the knockdown effects, which often exhibited differential effects of the two co-activators, whereby TAZ had a more robust effect on matrix degradation than YAP, The overexpression of each of them alone (and, certainly a combination of the two), had a comparable suppressive effect. This observation is consistent with the view that both YAP and TAZ suppress invadopodia, and that the differential effect of their knock down reflects variations in their relative prominence in the tested cell lines or their differential regulation by the Hippo core phosphorylation cascade (Shreberk-Shaked, Dassa et al. 2020).

The search for specific downstream targets, affected by YAP and TAZ knockdown included, primarily, proteomic and transcriptomic analysis of differentially expressed molecules (siYAP+TAZ vs. siControl), focusing on invadopodia-related components. Towards that goal, we have assembled a literature-curated invadopodia components database that contains structural and regulatory components of invadopodia and the associated cytoskeleton (Supplementary Table S7). Towards that end, we have identified the differentially expressed proteins and transcripts, detected by the proteomic and mRNA profiling, respectively. Overall, the proteomic data revealed significant changes in the levels of 150 proteins (72 were up regulated and 78 down regulated). Further, for each knockdown conditions, the differentially expressed proteins were found to be siYAP vs siControl (29 proteins) or siTAZ vs siControl (27 proteins), and siYAP+siTAZ vs siControl (94 proteins). In addition, some proteins (28) were common because they were identified as differentially expressed in more than one knockdown condition. Excluding the 28 proteins that were common in each knockdown condition results in 122 unique proteins in the proteomic profiling results obtained. The RNA seq data, showed differential expression of 675 transcripts (600 were upregulated and 75 down regulated). Further, for each knockdown conditions, the differentially expressed genes were found to be siYAP vs siControl (34 genes) or siTAZ vs siControl (61 genes), and siYAP+siTAZ vs siControl (580 genes). Notably, among the down-regulated molecules were known direct downstream transcriptional targets of the Hippo pathway, including YAP1 CTGF, CYR61. Comparing the proteomic and transcriptomic lists of differentially-expressed molecules revealed 15 overlapping components (F3, TGFB2, YAP1, SNAPC1, MARCKSL1, CYP1B1, TMEM163, PFKFB4, TNFSF10, GFPT2, NDRG1, GSN, ADAM8, ALDOC, RAB31; see Figure 7), among which there was just one molecule (gelsolin), which is included in our invadopodia list. Further data mining of the invadopodia related and differentially expressed proteins pointed to 7 proteins that were significantly elevated following YAP+TAZ knockdown, some of which are well known and essential invadopodia components. These include TKS5, an adaptor phospho-protein that is essential for invadopodia formation, recruitment of proteases and matrix degradation (25,27). Other upregulated molecules include MMP14, a key enzyme required for invadopodia-mediated matrix degradation (24), ADAM19, a member of the ADAM family (A Disintegrin And Metalloproteinase) (25,54), as well as the formin family member DIAPH2 (55), integrin □5, that, together with different □ chains form the adhesive domain of invadopodia (56,57), gelsolin, an actin modulator in invadopodia (58,59), that was also elevated transcriptionally, and IDH, whose specific function in invadopodia formation and matrix degradation is still unclear (39). Two invadopodia-related proteins that were downregulated following YAP+TAZ suppression were Serpine1 that can affect matrix degradation, and AKAP12, that affects PKA distribution in cells. The exact roles of these proteins in invadopodia-mediated caner invasion is unknown. Out of the 594 genes whose transcription was significantly affected by YAP/TAZ knockdown, 18 were present in our invadopodia-related list (Supplementary Table 10 and S11), including several integrin chains, matrix metallopeptidases, actin-associated proteins and signaling modulators.

Taken together these results suggest that invadopodia formation and matrix degradation activity are regulated at multiple levels, transcriptional and post-transcriptional. The proteomic and transcriptomic data presented here, clearly demonstrate that YAP and TAZ suppress essential structural components of invadopodia and suggest that Hippo-mediated inhibition of YAP/TAZ may increase the invasive phenotype. Further studies are warranted to better understand the precise role of the different Hippo pathway mediators that affect the activity of YAP and TAZ, either as individual proteins or as paralogs. Dissecting their contextual behavior and clarifying their conflicting functions in different cancer cell types will eventually enable the design and application of novel therapies targeting cancer invasion and metastasis.

## Acknowledgements

We thank the scientific staff at the Department of Life Sciences Core facilities, Weizmann Institute of Science (WIS), for the access to the research infrastructure used in this study, and for their competent help. We are particularly grateful to Dr. Yishai Levin at The De Botton Protein Profiling Institute of the Nancy and Stephen Grand Israel National Center for Personalized Medicine (G-INCPM), WIS, for the scientific input, guidance and help in the proteomic profiling. We are grateful to the Crown Genomics institute of the G-INCPM, WIS, for their assistance in RNA sequencing. We also thank Dr. Shlomit Reich-Zeliger (Prof. Nir Friedman Lab) for assistance with some of the reagents and access to instruments used for RNA quantitation and quality assessment of RNA samples for the RNA sequencing experiments in this study. This study was supported by grants from the Minerva Center for Aging, and a Precision Medicine grant from the Israel Science Foundation.

## Author contributions

J.B.V: Participated in the design and execution of the experiments included in this study preparation of the figures and in writing of the manuscript.

B.D Participated in the analysis of the transcriptomic and proteomic data, and in the assembly of the literature-based invadopodia proteome database used here.

D.M Participated in the proteomic profiling of the samples.

M.S.S. Participated in the experimental design of the YAP/TAZ modulation experiments. M.O. Participated in the experimental design and in the interpretation of the data.

B.G Participated in the design and follow-up of the experiments included in this study, data interpretation and in writing of the manuscript.

