## Supplementary Figures and Legends for "Deciphering the involvement of the Hippo pathway co-regulators, YAP/TAZ in invadopodia formation and matrix degradation"

**Supplementary Figure Legends**

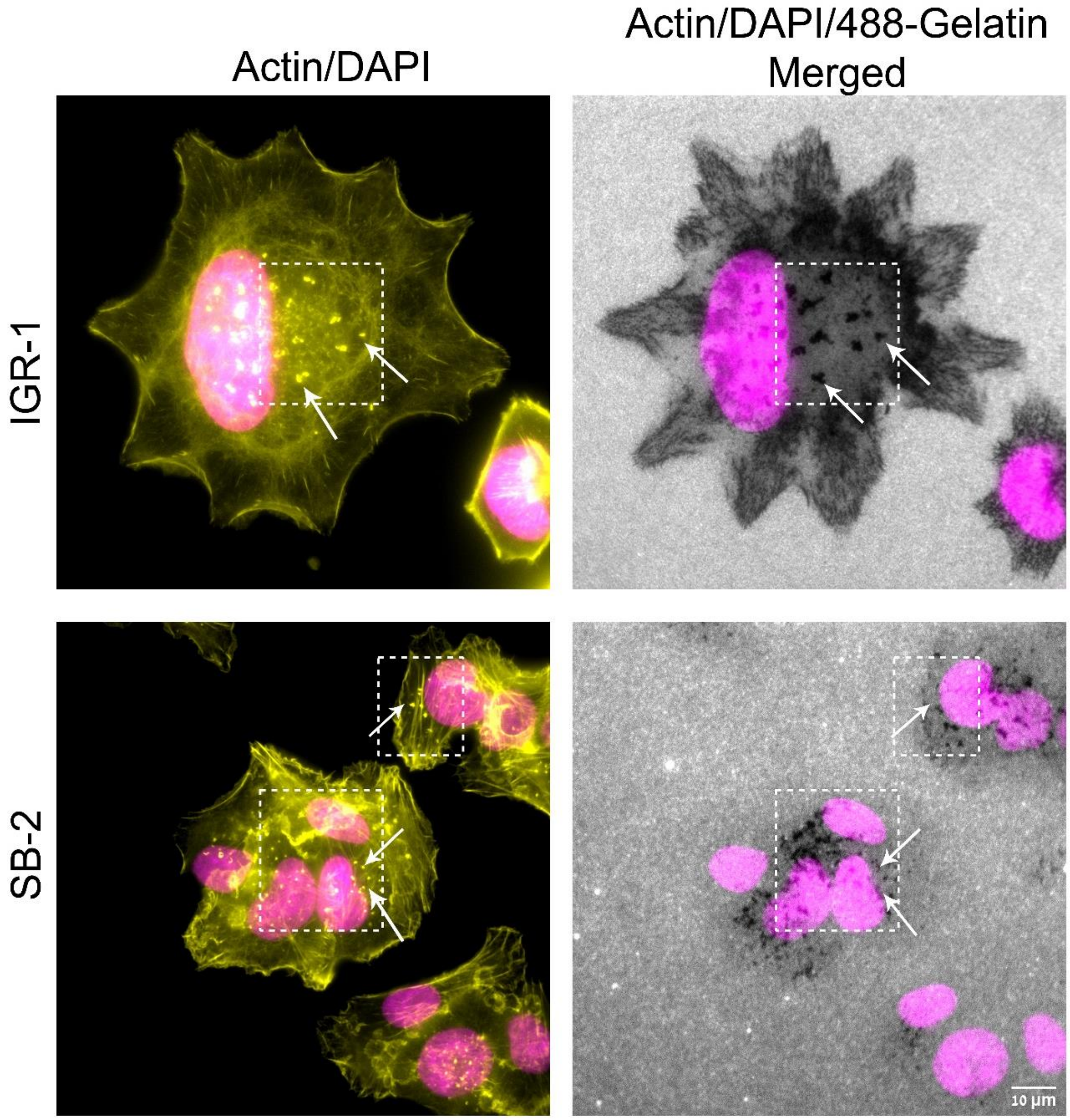


**Figure S1.** Zoomed-in images of cell lines, IGR-1 and SB-2 displaying invadopodia and their corresponding gelatin degradation. Cells were plated on Alexa 488-gelatin matrix for 5-6 h, subsequently fixed, and stained for actin and DAPI. Images were captured under 60 X (1.42) oil objective in a DeltaVision Elite microscopy system (Applied Precision, Issaquah, WA). Scale bar is 10 µm. A representative panel from four independent sets of experiments is shown here. Dotted line boxes (white) highlight some of the area where invadopodia actin dots (yellow) are present. Nuclei (magenta), and local gelatin matrix degradation visualized as dark spots on Alexa 488-gelatin are merged and shown here. Arrows (white) point out some of the invadopodia and its corresponding degradation on the Alexa 488-gelatin matrix.

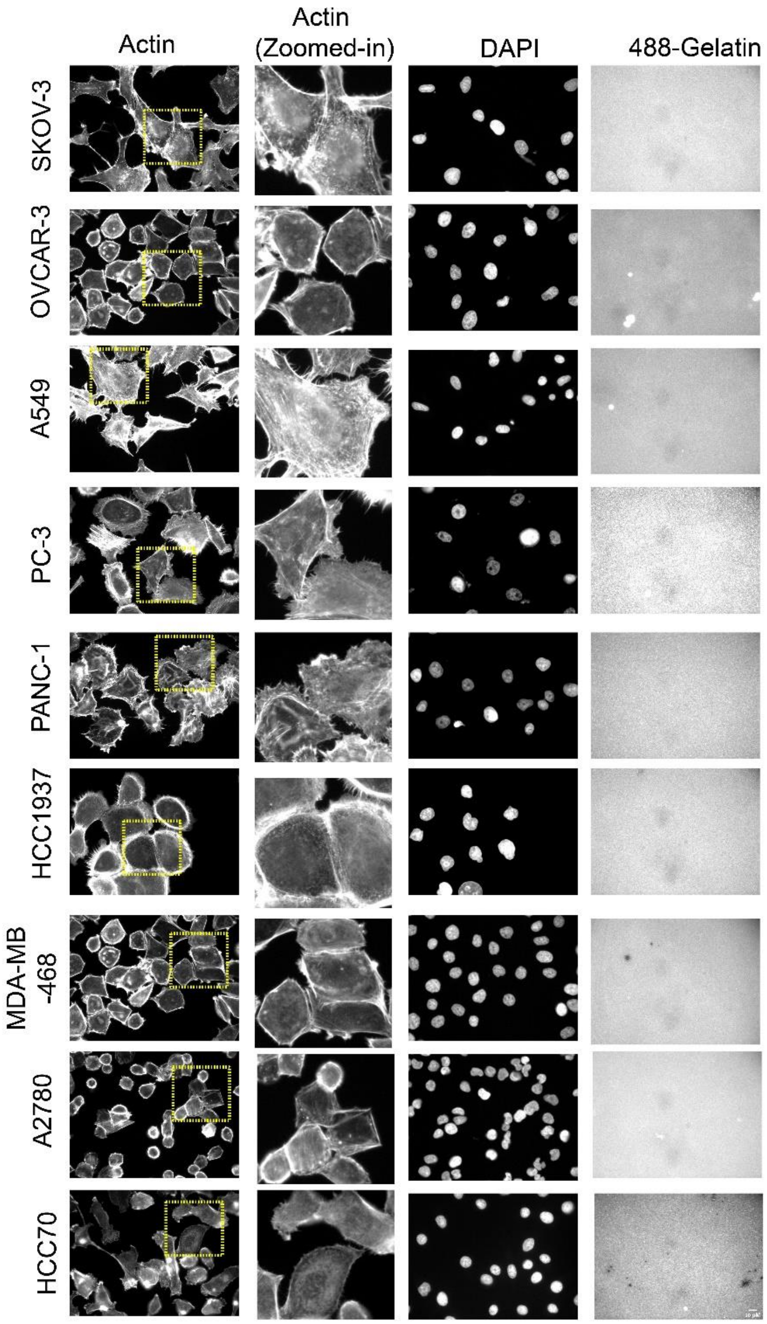


**Figure S2.** Multi-cancer cell line panel displaying gelatin degradation (< 1 μm$^2$/cell). Cancer cell lines SKOV-3, OVCAR-3, A549, PC3, PANC-1, HCC1937, MDA-MB-468, A2780, and HCC70 were plated on Alexa 488-gelatin and incubated for 5-6 h. Cells were fixed and stained for actin and DAPI. Images were captured under 40X (0.75) air objective using a WiScan® Hermes Imaging System (IDEA Bio-Medical Ltd.). Four independent sets of experiments were performed and a representative panel is shown here. A section of the cells in the actin panel is highlighted with dotted line yellow boxes to show zoomed-in images indicating the absence of invadopodia actin rich dots.

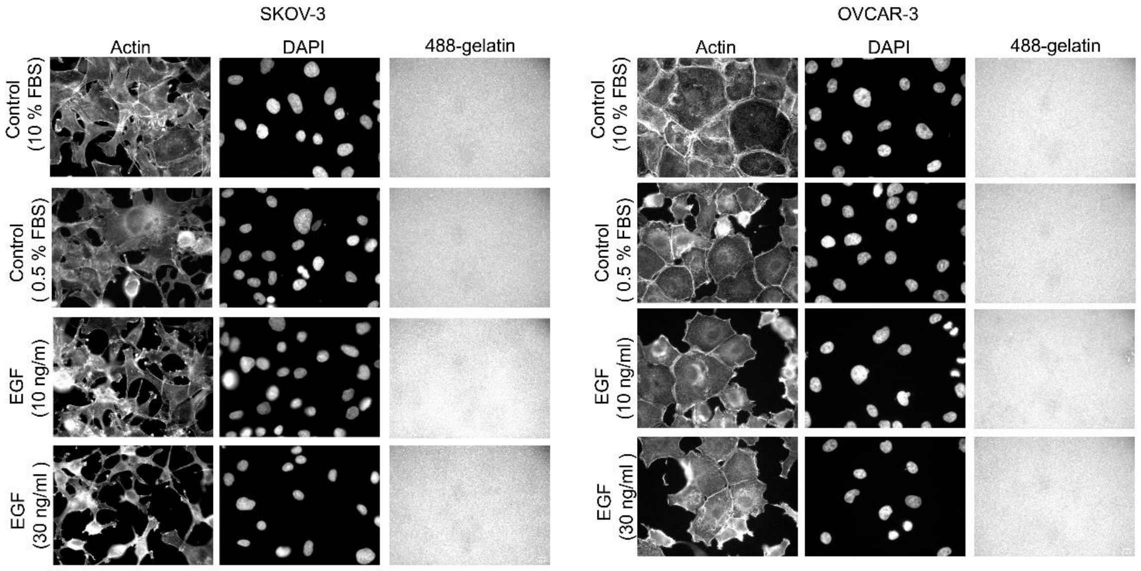


**Figure S3.** Effect of EGF stimulation on invadopodia formation and gelatin degradation observed in SKOV-3 and OVCAR-3 cancer cell lines. Cells were grown in medium containing 10 % FBS and serum starved (0.5 % FBS) for 24 h. Subsequently, cells grown in 0.5 % FBS and 10 % FBS were plated on Alexa 488-gelatin. Cells grown in 0.5 % FBS were stimulated by EGF (10 and 30 ng/ml) and incubated on Alexa 488-gelatin for 5-6 h. Cells were seeded on duplicate wells for each condition and 36 fields were obtained for each well. Images were captured under 40X (0.75) air objective using a WiScan® Hermes Imaging System (IDEA Bio-Medical Ltd.).

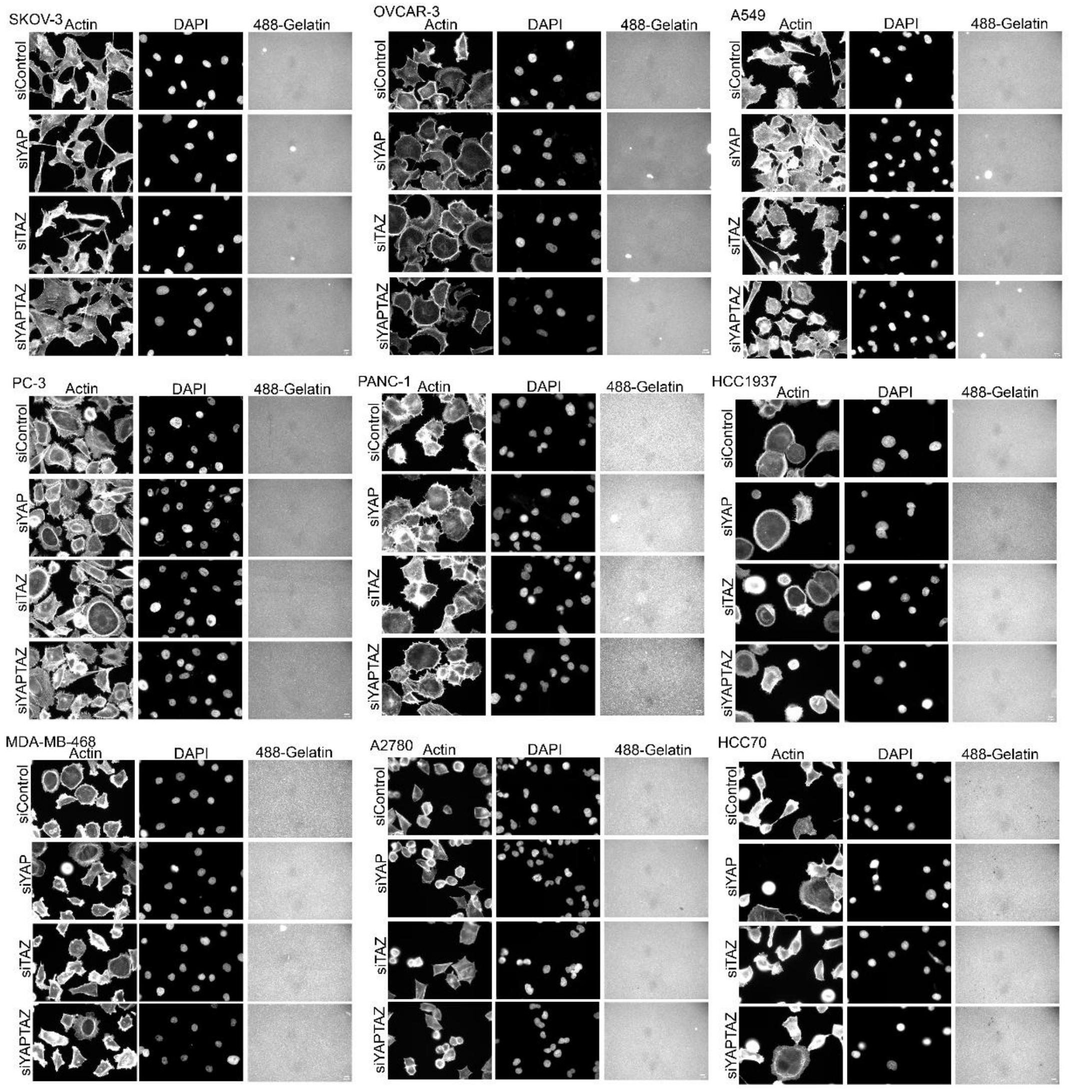


**Figure S4.** Nine cancer cell lines showing no effect on the invadopodia formation and subsequent gelatin degradation after the suppression of hippo co-regulators YAP, TAZ or both. Cancer cell lines SKOV-3, OVCAR-3, A549, PC3, PANC-1, HCC1937, MDA-MB-468, A2780, and HCC70 were knockdown for YAP, TAZ or both and subsequently plated on Alexa 488-gelatin for 5-6 h. Cells were fixed and stained for actin, DAPI and images were captured under 40X (0.75) air objective using a WiScan® Hermes Imaging System (IDEA Bio-Medical Ltd.). Cells were seeded on duplicate wells for each condition and images for 36 fields were obtained for each well. A representative panel from one of the set of experiment is shown here.

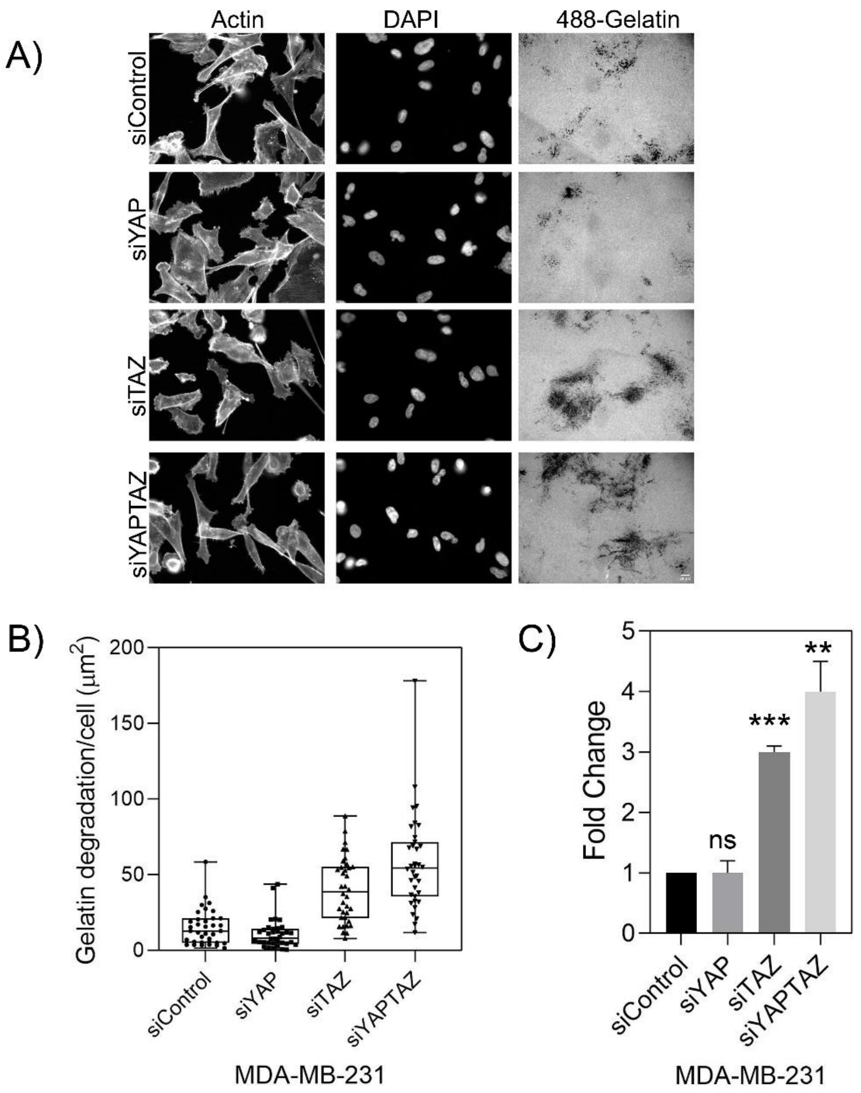


**Figure S5**. Knockdown of YAP, TAZ or YAP+TAZ using single oligonucleotides in MDA-MB-231 cell line is shown. Z-stack of images for actin, DAPI and Alexa Fluor 488-gelatin were captured under 40 X (0.75) air objective using a WIScan Hermes® microscope (IDEA Bio-Medical Ltd.). Cells were seeded on duplicate wells for each condition and 36 fields were obtained for each well. Four independent sets of experiment were performed. A). A representative panel of images are shown. Scale bar is 10 µm. B). An average plot of gelatin degradation/cell (µm$^2$) with ± SEM for each treatment condition is shown. C). Fold change obtained from the values for gelatin degradation/cell (µm$^2$) is plotted for each treatment condition. The significance values for the treatment conditions was calculated by paired t test. **, *P* value <0.0077, ***, *P* value <0.0001 ns, not significant. P value was <0.0052 (**) as determined by one-way ANOVA.

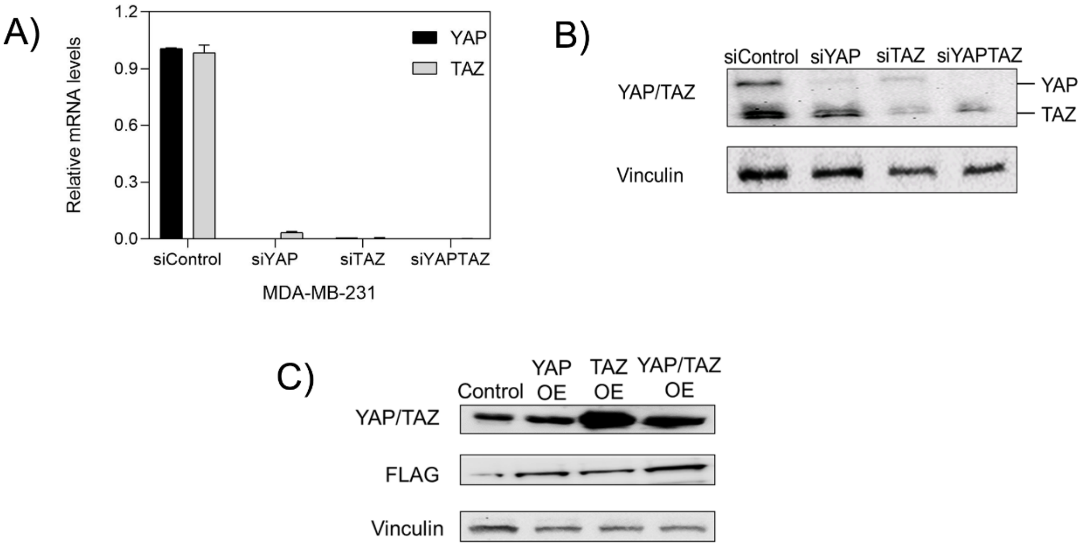


**Figure S**6. Validation of knockdown and overexpression of YAP/TAZ in MDA-MB-231 cells. A). RT-qPCR analysis of knockdown efficiency for siYAP, siTAZ and siYAP+TAZ mRNA levels in MDA-MB-231 cell line is shown. Data shown here is average of three independent set of experiments. (Mean ±SEM) $\log_2$ mRNA expression was normalized with respect to GAPDH/HPRT1 and siControl. The significance values for the treatment conditions by one way ANOVA was calculated as **, $P<0.0057$ for the shown treatment. B) Western blot of the knockdowns for each treatment condition is shown. C). Western blot after overexpression of YAP, TAZ and YAP +TAZ for 48 h.

A)

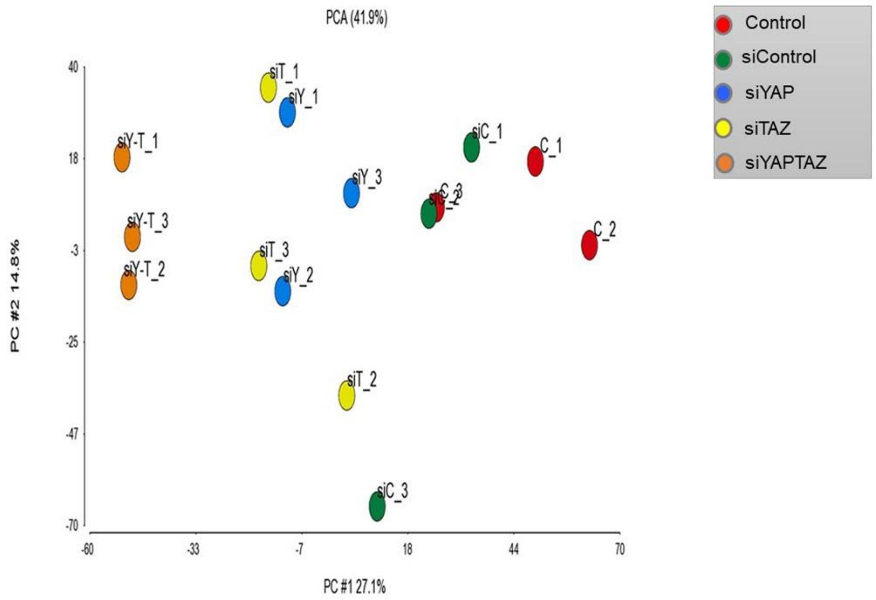


B)

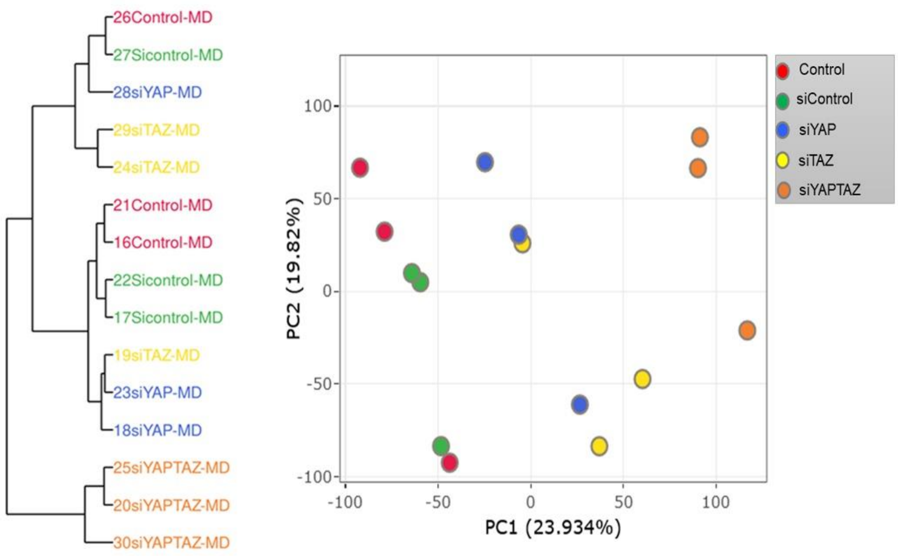


**Figure S7**. Principal component analysis (PCA) of the proteomics and transcriptomic profiling datasets upon knockdown of YAP, TAZ and YAP+TAZ in MDAMB-231 cell line. A). Principle component (PC) 1 demonstrates the split between the genotype, where the co-knockdown of YAP+TAZ is separated from the single knockdowns of YAP or TAZ, and from the controls. B). Principle component (PC1) of the RNA sequencing results for the co-knockdown of YAP+TAZ is separated from the single knockdowns of YAP or TAZ, and from the controls.

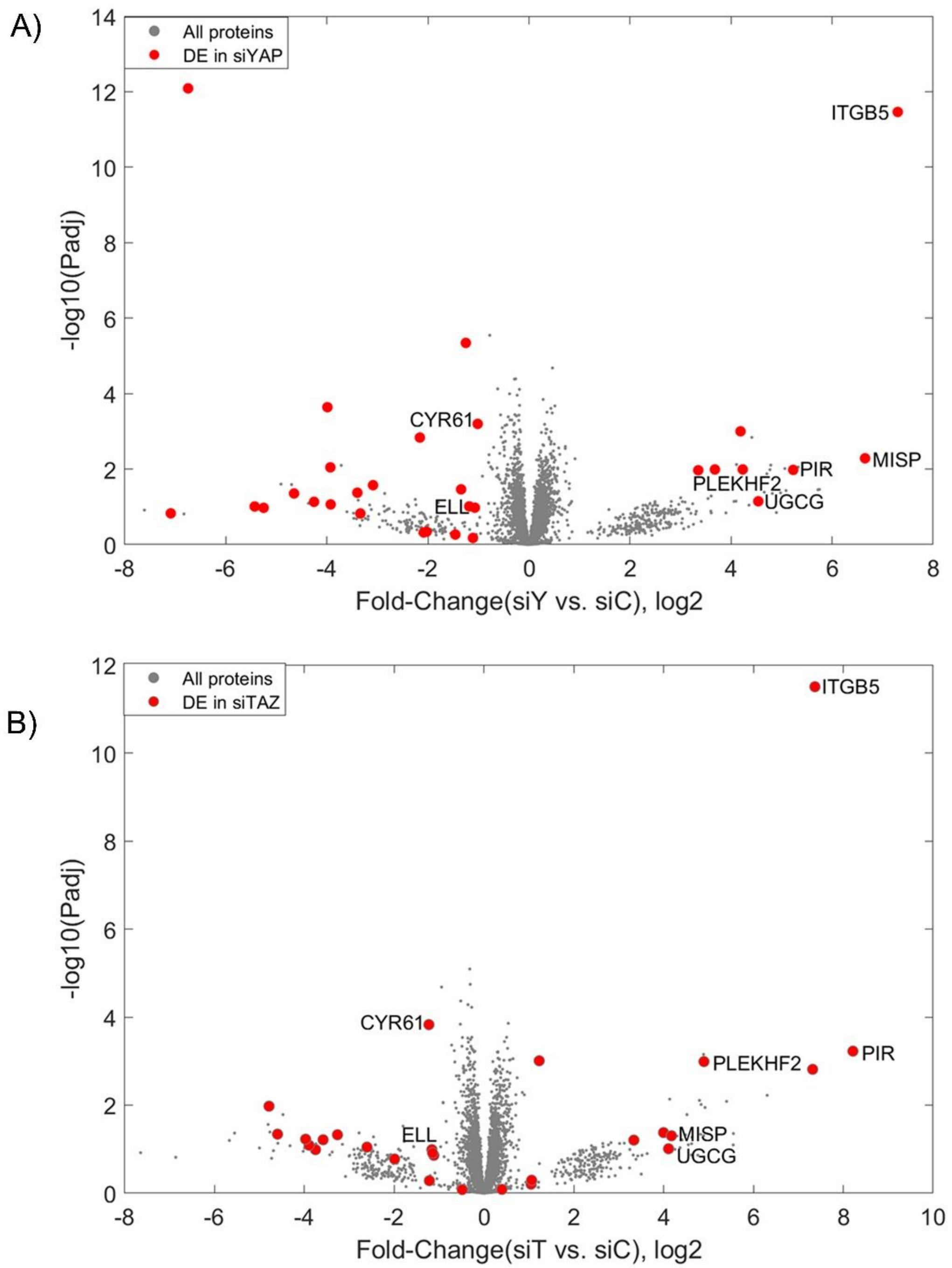


**Figure S8.** Volcano plots for the proteins detected after single knockdown of YAP or TAZ in MDA-MB-231 cells. A). All proteins that were detected upon single knockdown of YAP are shown in grey and differentially expressed proteins are shown in red. B). All proteins that were differentially expressed upon single knockdown of TAZ are shown in grey and differentially expressed proteins are shown in red. Proteins that were common and significantly expressed in both treatment conditions are labeled with their protein names.

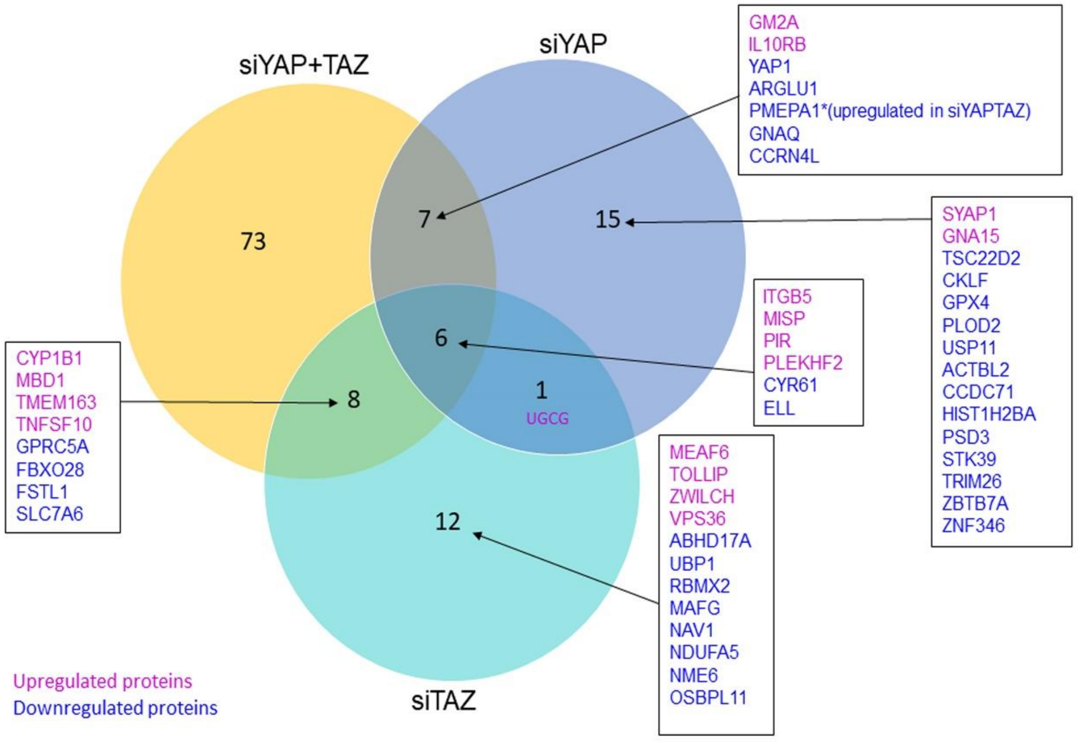


**Figure S9.** Venn diagram depicting the overlap between proteins differentially expressed in each treatment; siYAP, siTAZ and siYAP+TAZ are shown. Upregulated proteins are shown in magenta and down regulated proteins are shown in blue. Protein names are listed in boxes and in Supplementary Table S6.

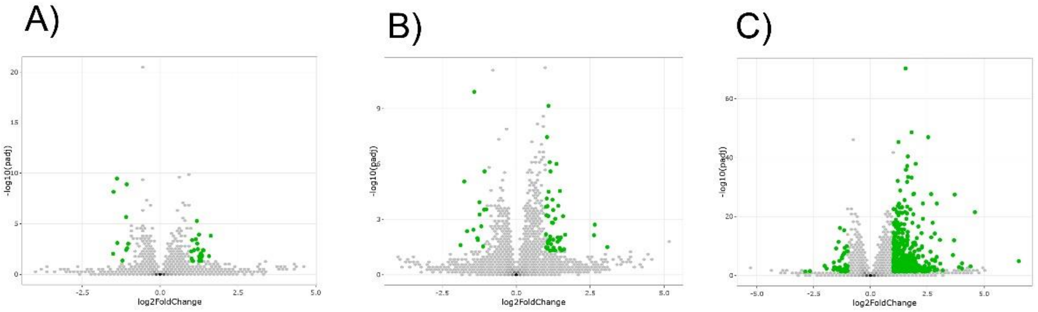


**Figure S10.** Volcano plots of differentially expressed genes after co-knockdown of siControl vs siYAP (34 genes), siControl vs siTAZ (61 genes) and siControl vs siYAPTAZ (580 genes) in MDA-MB-231 cell line are shown. The detailed breakdown of differentially expressed genes are listed in Supplementary Table S10.

**Supplementary Table Legends**

**Table S1**. List and characteristics of the multi-cancer cell line panel used in the study.
**Table S2**. List of SMARTpool oligonucleotides used in the study.
**Table S3**. List of single oligonucleotides used in the study.
**Table S4**. List of primers sequences used in the study.
**Table S5**. Proteomic profiling results (LFQ intensities) of all 4667 detected proteins in MDA-MB-231 cell line for each treatment conditions (Control, siControl, siYAP, siTAZ, siYAP+TAZ) obtained for three independent set of experiments with their ANOVA analysis and resultant fold change values are shown.
**Table S6**. Differentially expressed proteins analyzed from the protein profiling results for Control vs siControl, siControl vs siYAP, siControl vs siTAZ and siControl vs siYAPTAZ in MDA-MB-231 cell line for each independent set of experiment with their respective fold change and p values are enlisted.
**Table S7.** Curated list of proteins, structurally and functionally related to invadopodia obtained from published literature are listed.
**Table S8.** List of nine significantly upregulated and down-regulated proteins obtained after co-knockdown of YAP +TAZ in MDA-MB-231 cell line.
**Table S9.** RNA sequencing analysis for each treatment conditions (Control, siControl, siYAP, siTAZ, siYAP+TAZ) obtained for three independent set of experiments in MDA-MB-231 cell line is shown.
**Table S10**. List of invadopodia-associated genes in RNA-seq results upon knockdown of YAP, TAZ and YAP+TAZ in MDA-MB-231 cell line.
