## Supplementary material for "Deciphering the involvement of the Hippo pathway co-regulators, YAP/TAZ in invadopodia formation and matrix degradation": Table S1-S4

**Table S1.** List and characteristics of the multi-cancer cell line panel used in the study.

| Sr. No | Cell lines | Tissue/Disease | Invadopodia formation tested | Primary vs metastasis Source: CCLE/Published |
|---|---|---|---|---|
| 1 | MDA-MB-231 | Breast/adenocarcinoma | + | Metastasis |
| 2 | NCI-H1299 | Lung/non-small cell lung carcinoma | + | Metastasis |
| 3 | A-375 | Skin/malignant melanoma | + | Primary |
| 4 | CSK-A375* | Skin/malignant melanoma | + | Derived from A-375 |
| 5 | A2058 | Skin/melanoma | + | Metastasis |
| 6 | WM793 | Skin/melanoma | + | Primary |
| 7 | 63-T | Skin/melanoma | + | Metastasis |
| 8 | IGR-1 | Skin/melanoma | + | Metastasis |
| 9 | Malme-3M | Skin/ malignant melanoma | + | Metastasis |
| 10 | SB-2 | Melanocytes/malignant melanoma | + | Primary |
| 11 | UM-SCC-47 | Tongue/head and neck squamous cell carcinoma | + | Primary |
| 12 | LOX-IMVI | Skin/malignant melanoma | + | Metastasis |
| 13 | SKOV-3 | Ovary /adenocarcinoma | - | Metastasis |
| 14 | OVCAR-3 | Ovary /adenocarcinoma | - | Metastasis |
| 15 | A549 | Lung /carcinoma | - | Primary |
| 16 | PC-3 | Prostate /adenocarcinoma | - | Metastasis |
| 17 | PANC-1 | Pancreatic /carcinoma | - | Primary |
| 18 | HCC1937 | Breast /ductal carcinoma | - | Primary |
| 19 | MDA-MB-468 | Breast /adenocarcinoma | - | Metastasis |
| 20 | A2780 | Ovary /carcinoma | - | Primary |
| 21 | HCC70 | Breast/ductal carcinoma | - | Primary |

**Footnotes**

*CSK-A375: Knockdown of C-terminal Src kinase using shRNA in A375 cell line.

**Table S2.** List of SMARTpool oligonucleotides used in the study

| **Sr. No** | **siGenome** | **SMARTpool sequence** |
|---|---|---|
| 1 | Negative Control (Human) | UGGUUUACAUGUCGACUAA |
| 2 | YAP 1 (Human) | GCACCUAUCACUCUCGAGA;<br>GAACAUAGAAGGAGAGGAG;<br>CCACCAAGCUAGAUAAAGA;<br>GGUCAGAGAUACUUCUUAA |
| 3 | TAZ (Human) | AAGCCUAGCUCGUGGCGGA;<br>AGGAACAAACGUUGACUUA;<br>GGACAAACACCCAUGAACA;<br>GACAUGAGAUCCAUCACUA |

**Table S3.** List of single oligonucleotides used in the study

| **Sr. No** | **siGenome** | **Single oligo sequence** |
|---|---|---|
| 1 | YAP 1 (Human) | 1) CCACCAAGCUAGAUAAAGA<br>2) GCACCUAUCACUCUCGAGA |
| 2 | TAZ (Human) | 1) GGACAAACACCCAUGAACA<br>2) AAGCCUAGCUCGUGGCGGA |

**Table S4.** List of primers sequences used in the study

| Sr. No | Gene Name | Primer sequence |
|---|---|---|
| 1 | YAP1 (Human) | Forward-GCCGGAGCCCAAATCC<br>Reverse-GCAGAGAAGCTGGAGAGGAATG |
| 2 | TAZ (Human) | Forward-CGATGACCCCAGACATGAGA<br>Reverse-CTCGAATGATATGGCCCTCC |
| 3 | GAPDH | Forward- ACCCACTCCTCCACCTTTGA<br>Reverse- CTGTTGCTGTAGCCAAATTCGT |
| 4 | HPRT | Forward- CTGAGGATTTGGAAAGGGTGT<br>Reverse- CATCTCGAGCAAGACGTTCA |

**Supplementary Table Legends**

**Table S1**. List and characteristics of the multi-cancer cell line panel used in the study.
**Table S2**. List of SMARTpool oligonucleotides used in the study.
**Table S3**. List of single oligonucleotides used in the study.
**Table S4**. List of primers sequences used in the study.
**Table S5**. Proteomic profiling results (LFQ intensities) of all 4667 detected proteins in MDA-MB-231 cell line for each treatment conditions (Control, siControl, siYAP, siTAZ, siYAP+TAZ) obtained for three independent set of experiments with their ANOVA analysis and resultant fold change values are shown.
**Table S6**. Differentially expressed proteins analyzed from the protein profiling results for Control vs siControl, siControl vs siYAP, siControl vs siTAZ and siControl vs siYAPTAZ in MDA-MB-231 cell line for each independent set of experiment with their respective fold change and p values are enlisted.
**Table S7.** Curated list of proteins, structurally and functionally related to invadopodia obtained from published literature are listed.
**Table S8.** List of nine significantly upregulated and down-regulated proteins obtained after co-knockdown of YAP +TAZ in MDA-MB-231 cell line.
**Table S9.** RNA sequencing analysis for each treatment conditions (Control, siControl, siYAP, siTAZ, siYAP+TAZ) obtained for three independent set of experiments in MDA-MB-231 cell line is shown.
**Table S10**. List of 18 invadopodia-associated genes found to be differential in RNA-seq results upon co-knockdown of YAP+TAZ in MDA-MB-231 cell line.
**Table S11.** List of invadopodia-associated (highlighted in yellow) or hippo pathway related genes (highlighted in bold) obtained in RNA-seq results upon knockdown of YAP, TAZ and YAP+TAZ in MDA-MB-231 cell line.
